# An Omic and Multidimensional Spatial Atlas from Serial Biopsies of an Evolving Metastatic Breast Cancer

**DOI:** 10.1101/2020.12.03.408500

**Authors:** Brett E. Johnson, Allison L. Creason, Jayne M. Stommel, Jamie M. Keck, Swapnil Parmar, Courtney B. Betts, Aurora Blucher, Christopher Boniface, Elmar Bucher, Erik Burlingame, Todd Camp, Koei Chin, Jennifer Eng, Joseph Estabrook, Heidi S. Feiler, Zhi Hu, Annette Kolodzie, Ben L. Kong, Marilyne Labrie, Jinho Lee, Patrick Leyshock, Souraya Mitri, Janice Patterson, Jessica L. Riesterer, Shamilene Sivagnanam, Julia Somers, Damir Sudar, Guillaume Thibault, Christina Zheng, Xiaolin Nan, Laura M. Heiser, Paul T. Spellman, George Thomas, Emek Demir, Young Hwan Chang, Lisa M. Coussens, Alexander R. Guimaraes, Christopher Corless, Jeremy Goecks, Raymond Bergan, Zahi Mitri, Gordon B. Mills, Joe W. Gray

## Abstract

Mechanisms of therapeutic resistance manifest in metastatic cancers as tumor cell intrinsic alterations and extrinsic microenvironmental influences that can change during treatment. To support the development of methods for the identification of these mechanisms in individual patients, we present here an Omic and Multidimensional Spatial (OMS) Atlas generated from four serial biopsies of a metastatic breast cancer patient during 3.5 years of therapy. This resource links detailed, longitudinal clinical metadata including treatment times and doses, anatomic imaging, and blood-based response measurements to exploratory analytics including comprehensive DNA, RNA, and protein profiles, images of multiplexed immunostaining, and 2- and 3-dimensional scanning electron micrographs. These data reveal aspects of therapy-associated heterogeneity and evolution of the cancer’s genome, signaling pathways, immune microenvironment, cellular composition and organization, and ultrastructure. We present illustrative examples showing how integrative analyses of these data provide insights into potential mechanisms of response and resistance, and suggest novel therapeutic vulnerabilities.

## Introduction

Precision Medicine is an approach to disease management that selects treatments based on the presence of one or more molecular, environmental, or lifestyle features that are associated with a positive therapeutic response. Applied to cancer, this approach has led to substantial improvements in clinical outcomes, increasingly through the use of analytical procedures to identify patients with molecular characteristics whose presence is associated with an increased likelihood of responding.^1,2^ These biomarkers can include genomic or proteomic abnormalities that activate signaling pathways on which cancers depend for survival, molecular features that define regulatory networks that control therapeutically-vulnerable cancer “hallmarks”, microenvironmental architectures, or aspects of immune dysfunctions.

Unfortunately, treatments deployed according to precision medicine principles do not always elicit positive responses and durable control is achieved for only a subset of patients with metastatic cancer.^3^ We posit that the failure to control individual cancers using biomarker-guided treatments stems in large part from our imperfect understanding of the multitude of critical characteristics that drive individual tumors and their adaptive ability to survive. These mechanisms of resistance may involve regulatory networks intrinsic to tumor cells, chemical and mechanical signals from proximal or distal microenvironments, and/or aspects of the immune system. They may vary between patients with similar guiding biomarkers, across metastases within a patient, and among tumor cell subpopulations within a single lesion, and they may also change during treatment.

To support development of methods to identify diverse mechanisms that influence progression and responses to treatment in individual patients, we present here a comprehensive Omic and Multidimensional Spatial (OMS) Atlas describing treatments, anatomic images, longitudinal blood biomarkers, and omic and image analyses of serial biopsies from a single patient with metastatic breast cancer. We applied a diverse array of molecular, microscopic, and quantitative analysis techniques to this patient’s primary tumor and four serial metastatic biopsies taken during 3.5 years of treatment. The combined analytic results describe the cellular and molecular composition and organization of the multiple biopsies and include: 1) comprehensive DNA, RNA, and protein profiles; 2) molecular signaling pathways and transcriptional regulatory networks; 3) immune, stromal, and tumor cell composition and organization; and 4) 2D and 3D subcellular and extracellular ultrastructure. The resulting data and integrative analyses were combined with detailed information about treatments and responses as revealed by frequent measurements of tumor biomarkers in blood as well as by computed tomography (CT) and fluorodeoxyglucose-positron emission tomography (FDG-PET) imaging. Both the clinical and biological studies were carried out under the Serial Measurements of Molecular and Architectural Responses to Therapy (SMMART) program at Oregon Health & Science University (OHSU), with support from the Human Tumor Atlas Network (HTAN).^4,5^

In the following sections, we describe the clinical and experimental workflows used to acquire and manage the data as well as the resulting datasets that comprise the OMS Atlas, which are available through the HTAN Data Coordinating Center (https://humantumoratlas.org/). We also present several integrative computational analyses to illustrate the utility of the Atlas in exploring spatial and temporal heterogeneity, understanding tumor evolution, elucidating resistance mechanisms, and revealing novel therapeutic vulnerabilities.

## Results

### Workflows for the implementation of personalized medicine

We established the SMMART program and HTAN workflows for rapid collection and interpretation of information about treatments, tumor responses, and the molecular and cellular characteristics of tumor lesions in ways that can support clinical decision making and mechanistic discovery (Figure 1). All patients in this program consent to participate in the IRB-approved observational study Molecular Mechanisms of Tumor Evolution and Resistance to Therapy (MMTERT). Under the MMTERT protocol, patients are monitored via periodic CT and FDG-PET imaging to quantify the sizes and metabolic activities of individual metastatic lesions. Blood analyses are also performed that report on tumor protein biomarkers, circulating tumor DNA (ctDNA) levels and genetic alterations, and standard blood compositions. Further, serial core needle biopsies of selected metastatic lesions provide tumor tissue that is divided and preserved for downstream assays within two to five minutes of removal from the patient for optimal preservation of molecular and architectural features. Biospecimens are used for both clinical analytics, which are performed in a CLIA-certified, CAP-accredited laboratory and are available to support clinical decision-making, and exploratory analytics, which are performed in academic research laboratories or core facilities. Manual and automated abstraction from the patient’s medical record generates a set of clinical metadata, including detailed information about anticancer treatments and supportive care, for integration with the analytic results.

**Figure 1.**
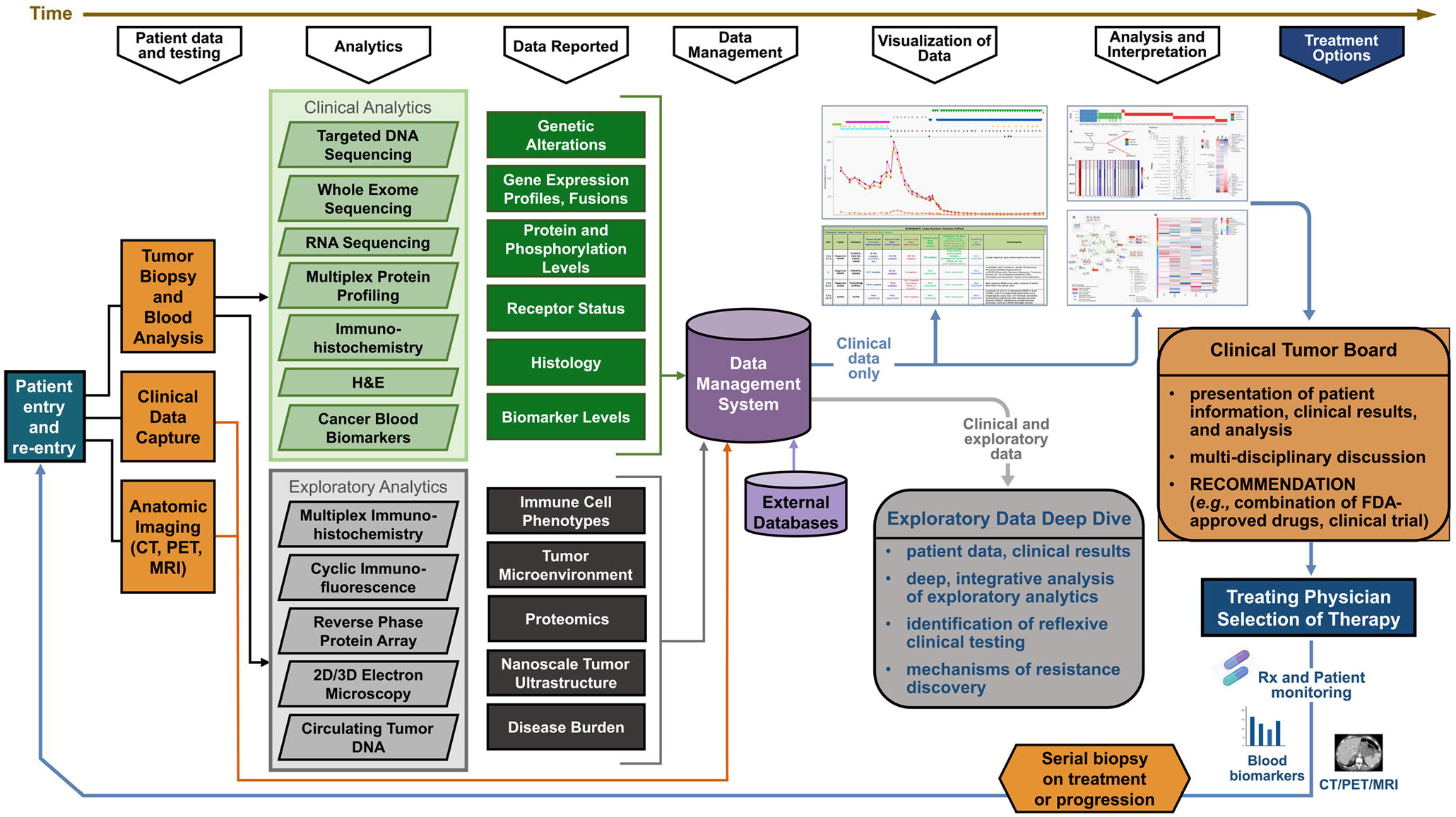
Workflows and analytical platforms used in generation of the OMS Atlas

We combine several software applications to create a robust computational platform for management of biospecimens as well as analysis and visualization of the resultant clinical and analytic data. Biospecimens are tracked and managed using a custom implementation of the LabVantage laboratory information management system. The LabKey system is used to store and visualize both clinical data and results from analysis workflows.^6^ The Galaxy computational workbench is used to create and run multi-step analysis workflows that process raw omics and imaging datasets.^7^ The OMERO system is used to visualize multiplex imaging and electron microscopy datasets and associated metadata.^8^ Ultimately, patient information and clinical assay results are presented to a multi-disciplinary clinical tumor board charged with providing personalized evidence-based recommendations to support combination therapies. Final treatment selection is at the discretion of the treating physician and patient. De-identified data are available to the research community to support mechanistic discovery efforts.

### Longitudinal data generation from a single metastatic breast cancer patient

The focus of this OMS Atlas is a female diagnosed with hormone receptor-positive, HER2 normal, right breast ductal carcinoma at the age of 64. She underwent a lumpectomy with intra-operative radiation therapy followed by treatment with four cycles of adjuvant docetaxel and cyclophosphamide chemotherapy. She then received two years of anastrozole treatment, followed by exemestane for five years. At that time, an ultrasound prompted by persistent abdominal discomfort revealed at least three liver masses, suggestive of metastasis. Staging CT scans revealed widespread metastatic disease involving mediastinal lymph nodes, including the hila bilaterally, lung parenchyma, liver, spleen, right adrenal gland, retroperitoneal lymph nodes, and likely skeleton. An FDG-PET bone scan confirmed metastatic bone disease. The patient was then enrolled in the SMMART program and consented to the MMTERT observational study.

Execution of the SMMART/HTAN workflow enabled the development of this OMS Atlas that captures the evolution of this metastatic breast cancer in response to treatment in four phases over a 3.5-year period (Figure 2A). Phase 1 treatment consisted of palbociclib, everolimus, and fulvestrant. Phase 2 consisted of doxorubicin and pembrolizumab. Phase 3 consisted of enzalutamide, capecitabine, fulvestrant, and the continuation of pembrolizumab and fulvestrant. Phase 4 consisted of carboplatin. Denosumab, pegfilgrastim, and hydroxychloroquine were given as supportive care. Temporary tumor control was achieved in the first three phases, with a new phase of therapy beginning upon signs of progression. Representative CT, FDG-PET, and ultrasound images highlight disease burden at key timepoints during the four phases of treatment (Figures S1A-L).

**Figure 2.**
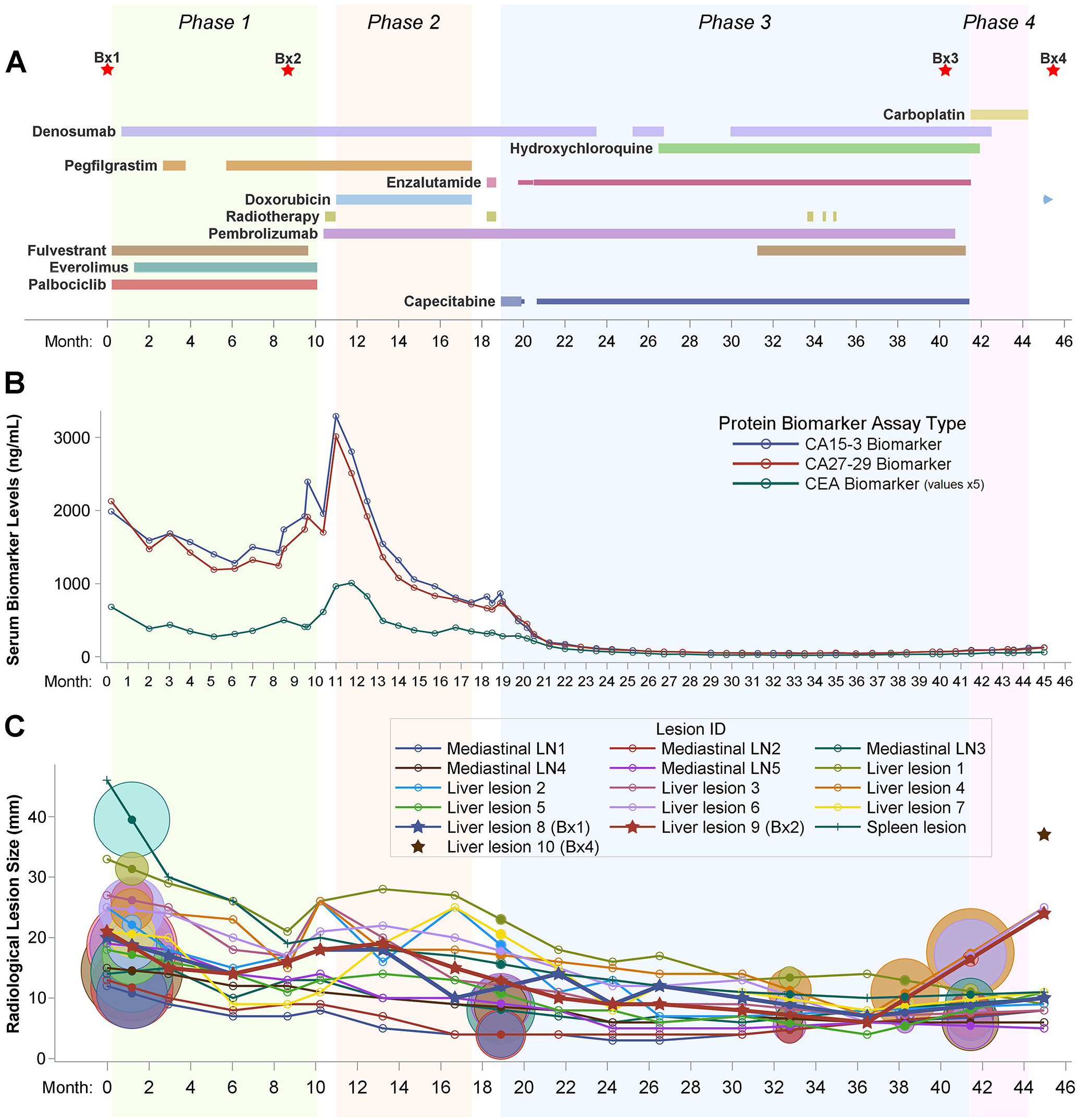
Timeline of clinical treatment and response metrics (A) Treatment schedule and biopsy timing (red stars) over the course of four phases of treatment (green, orange, blue, and pink areas). Timeline sectioned into 28-day months. The duration and relative dose for each drug is indicated by the extent and width of a horizontal bar, respectively. Continuation of a drug after the end of Phase 4 is indicated by a right pointing arrow. (B) Clinically reported serum levels of three tumor protein biomarkers. CEA values were multiplied by 5 to facilitate visualization. (C) Longitudinal tracking and variation in the longest-axis size of 16 representative metastatic lesions measured from serial CT images. Targets of metastatic biopsies are bolded and marked with stars. Circles represent FDG-PET imaging results, colored and centered on the lines of their corresponding lesion at interpolated lesion sizes. The diameter of each circle is proportional to the background-normalized maximum standardized uptake value (SUVmax). See also Figures S1, S2, and Table S1.

Abstracted clinical metadata in Table S1 links detailed treatment doses and timelines (Figures 2A, S2A) to tumor response metrics, including scheduled collection of CEA, CA 15-3, and CA 27-29 serum tumor protein biomarker levels (Figure 2B); the sizes and metabolic activities of representative metastatic lesions from periodic CT and FDG-PET scans (Figure 2C); as well as standard toxicity-related blood chemistries that include absolute neutrophil count, platelet, and liver function tests (Figures S2B-D).

Biospecimens collected include serial blood samples, a primary breast tumor, and four metastatic lesions: a liver biopsy taken immediately prior to Phase 1 (Bx1); a biopsy of a different liver lesion taken at the end of Phase 1 (Bx2); a bone lesion biopsy taken at the end of Phase 3 (Bx3); and a biopsy of a third liver lesion taken at the end of Phase 4 (Bx4; Figure 2A). Bx2, Bx3, and Bx4 were acquired from metastatic lesions explicitly identified on serial CT and/or FDG-PET imaging as progressing near the end of each respective treatment phase (Figures S1A-I). Importantly, the changes in size of each lesion before and after biopsy were recorded when visible by CT imaging (Figure 2C).

The OMS Atlas includes results from ten distinct omic and multiscale spatial imaging assays that were applied, when tissue availability allowed, to the primary tumor and four metastatic biopsies. Table S1 summarizes clinically reported results of immunohistochemistry (IHC) staining. Figures 3 and S3 highlight the differences in genomic alterations, gene expression, protein levels, and computed pathway activities of the biopsies with relevant comparisons to breast cancers from The Cancer Genome Atlas (TCGA).^9^ Figures 3A and 3B summarize the results of DNA analyses used to identify mutations and copy number alterations by targeted DNA sequencing of a clinically curated panel of cancer genes (GeneTrails^®^ Solid Tumor Panel), by whole exome sequencing (WES), and by low-pass whole genome sequencing (LP-WGS). Figures S2E and S3A summarize the results of whole exome and Dual Index Degenerate Adaptor Sequencing (DIDA-Seq) of ctDNA from serial blood samples.^10^ Figure 3 also summarizes the results from RNA sequencing (RNAseq) to characterize the whole transcriptome (Figure 3C), reverse phase protein arrays (RPPA) to profile the abundance of 450 proteins and phosphoproteins (Figures 3D, 3E), and a clinical multiplex protein analysis (Intracellular Signaling Protein Panel) of 23 proteins and phosphoproteins implicated in cancer (Figure 3F).^11–13^ Figures 4 and S4 show changes between the primary tumor and four metastases in the composition and functional status of individual leukocyte lineages using multiplex immunohistochemistry (mIHC).^14–16^ Figures 5 and S5 assess tumor and microenvironment cellular composition, functional state, and spatial organization using cyclic immunofluorescence (CycIF) and focused ion beam-scanning electron microscopy (FIB-SEM).^17–19^ Figures 6 and S6A show additional two-dimensional (2D) and three-dimensional (3D) cellular and subcellular features revealed using FIB-SEM.

**Figure 3.**
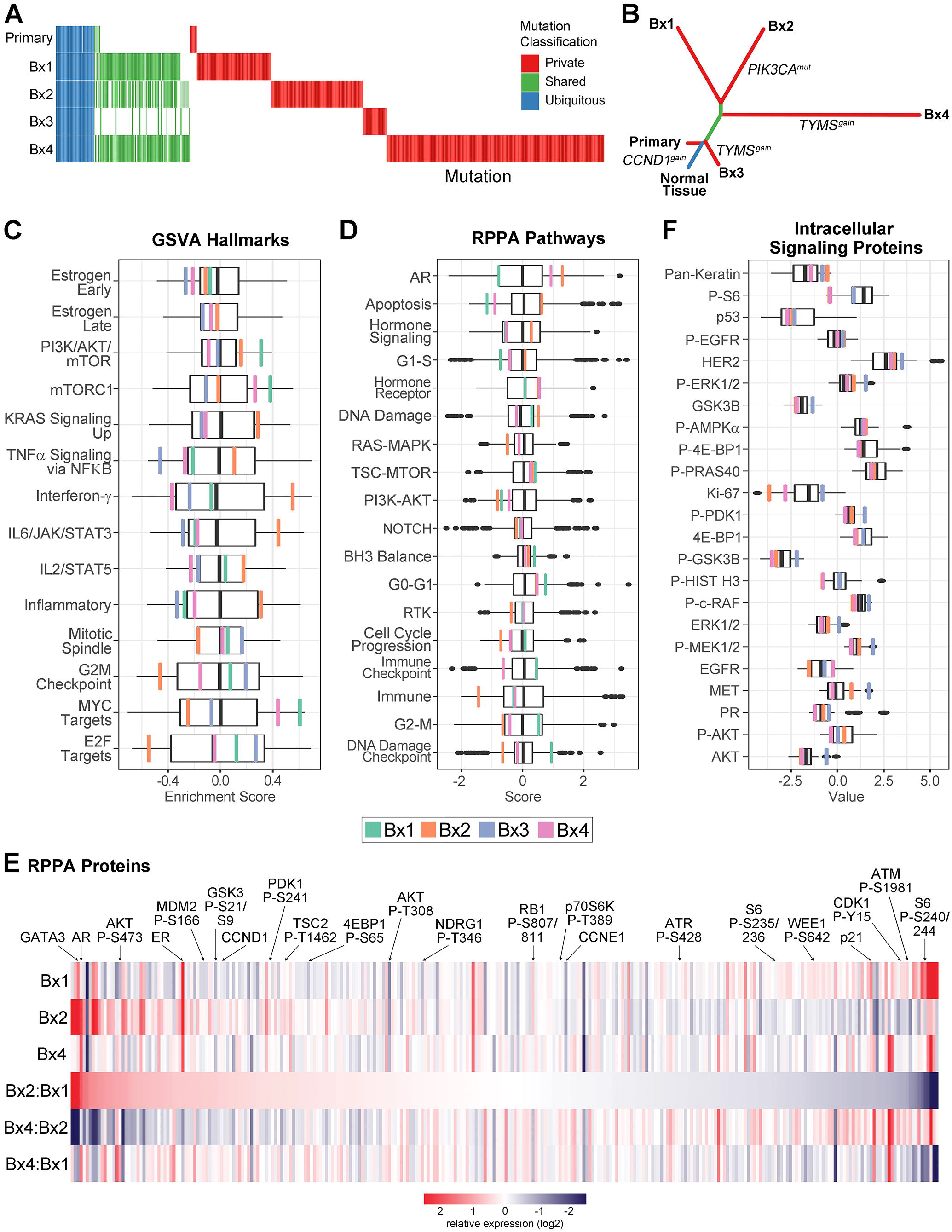
Genomic, transcriptomic, and proteomic profiling reveals spatiotemporal heterogeneity and evolution (A) Comparison of somatic mutations. Columns represent individual, non-silent SNVs or Indels identified from WES of at least one tissue sample and classified as Ubiquitous (present in all samples; blue), Shared (present in at least two samples; green), or Private (present in only a single sample; red). Mutational status in each sample is indicated as independently called (colored), detected in at least 2 sequencing reads but not independently called (reduced opacity), or absent (white). (B) Phylogenetic tree showing the evolutionary relationship between the primary tumor and four metastases. A *CCND1* gain was ubiquitous, a pathogenic *PIK3CA* p.E542K mutation was private to Bx2, and a *TYMS* amplification was shared by Bx3 and Bx4. (C) Transcriptomic gene set variation analysis (GSVA) of Cancer Hallmark pathways. The boxplot represents the distribution (upper and lower quartiles and median) of GSVA scores for the TCGA Luminal breast cancer cohort. Enrichment scores are shown for each of the biopsy samples: Bx1 (green), Bx2 (orange), and Bx3 (blue), and Bx4 (pink). (D) RPPA protein pathway activity assessment using pathway scores. The boxplots represent the distribution of the pathway activity of the TCGA breast cancer cohort. The pathway activities of three biopsy samples are marked as in D. (E) Total and phosphoprotein levels from RPPA normalized within the TCGA breast cancer cohort. The heat map shows relative protein levels for three biopsy samples and the fold-change between sample pairs. Proteins are ordered based on the fold-change difference between Bx2 relative to Bx1. Selected proteins are highlighted. (F) Intracellular Signaling Protein Panel measurements of total and phosphoprotein levels. The boxplots represent the distribution of protein levels of 57 metastatic breast cancers. The protein levels of three biopsy samples are marked as in D. See also Figure S3 and Table S2.

**Figure 4.**
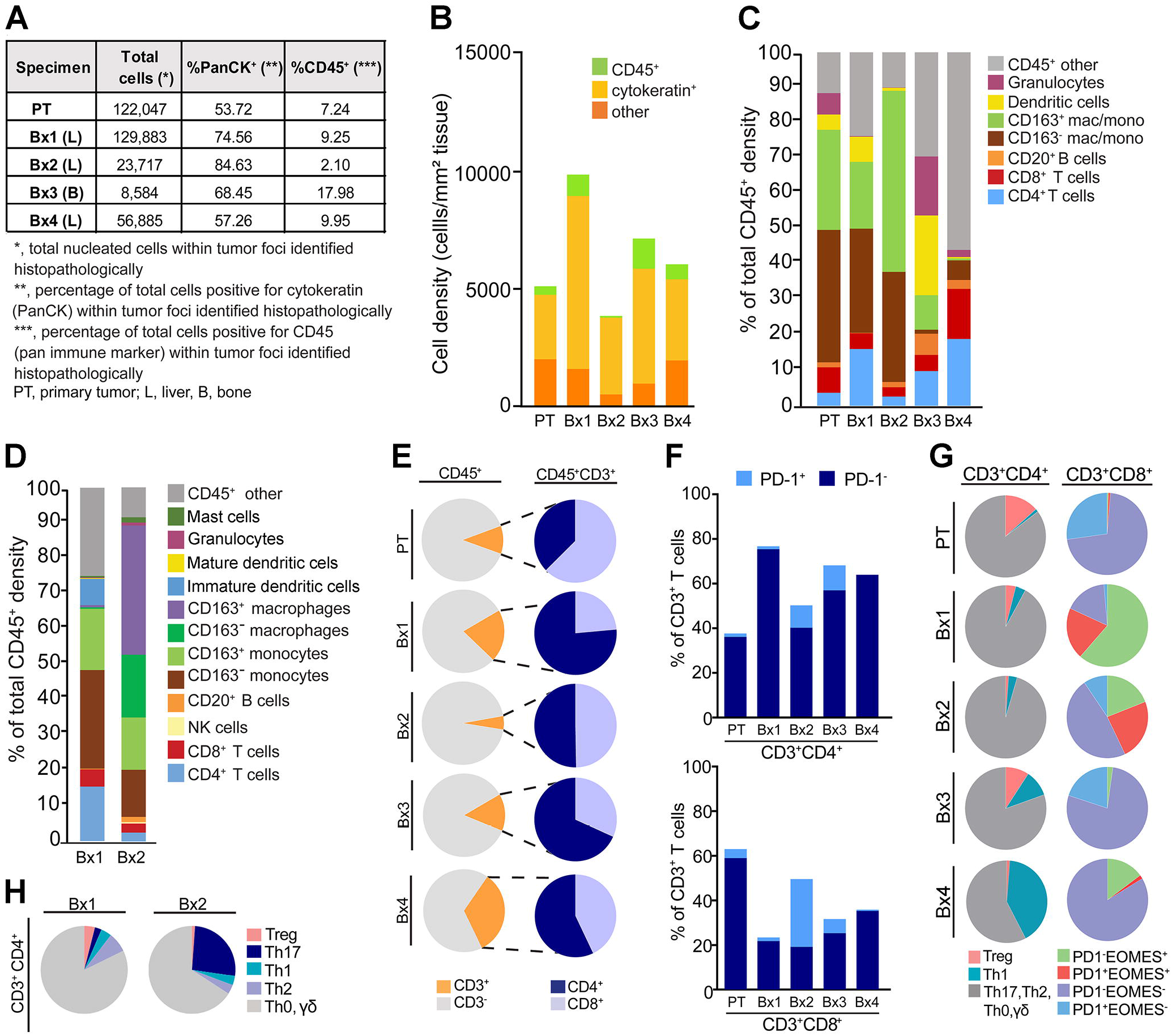
Monitoring response to therapy with deep *in situ* immune phenotyping by mIHC (A) Primary tumor (PT) and four biopsies (Bx1, Bx2, Bx3, and Bx4) were subjected to multiplex immunohistochemical (mIHC) analyses quantitatively evaluating immune (CD45^+^) and epithelial (Pan cytokeratin/CK^+^) positive cells in tumor compartments enumerated as percent of total nucleated cells. (B) Graphical representation of tissue composition, showing overall cell density (#cells/mm^2^ tissue analyzed) of PanCK^+^, CD45^+^, and PanCK^-^ CD45^-^ (other) nucleated cells. (C) Immune composition (as percent of total CD45^+^ cells) comparisons of seven major leukocyte lineages. (D) Deeper auditing of leukocyte lineages in Bx1 and Bx2 enumerating 12 immune cell populations and functional states. (E) Total CD3^+^ T cell abundance (orange pie slice) of total CD45^+^ cell populations (left), and proportion of CD4^+^ (blue) and CD8^+^ T cells (periwinkle) within total T cells (right). (F) Presence of PD-1^+^ cells (as percent of total CD3^+^T cells) in both the CD3^+^CD4^+^ (top) and CD3^+^CD8^+^ (bottom) T cell populations. (G) Differentiation state of CD3^+^CD4^+^ T cells reflected by regulatory (Treg), Th1, and Th2, Th17, Th0/γδ subsets (left), and CD3^+^CD8^+^ T cells as reflected by expression of PD-1 and EOMES. (H) Differentiation state of CD3^+^CD4^+^ T cells reflected by regulatory (Treg), Th17, Th1, Th2, and Th0/γδ subsets in Bx1 and Bx2. See also Figure S4 and Table S2.

**Figure 5.**
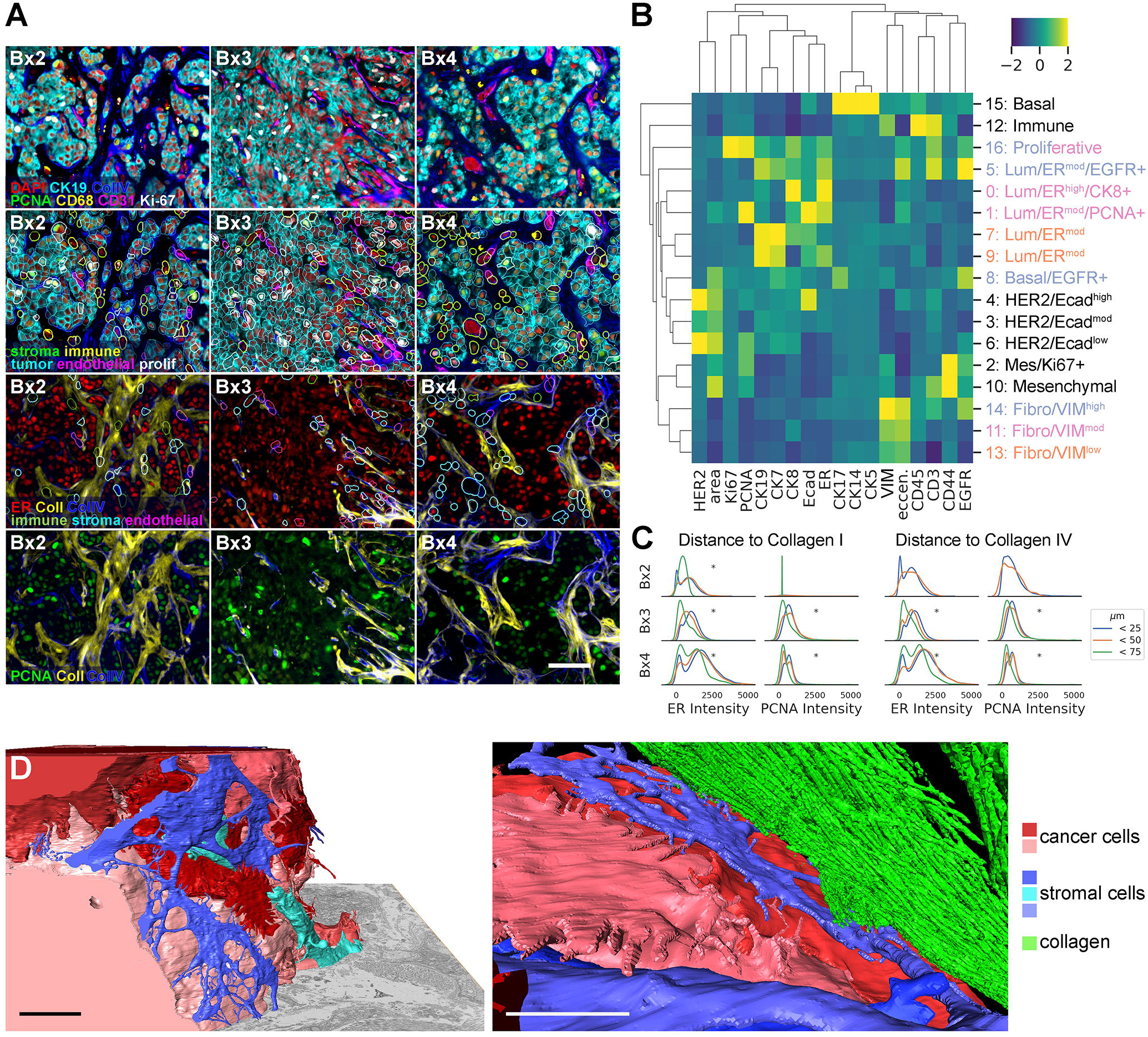
Monitoring tumor and stromal responses to therapy using CycIF and FIB-SEM (A) Example images of antibody staining overlaid with segmentation borders colored by cell type. See methods for gating details. Scale bar, 50 μm. (B) Heat map of mean z-scored intensity of unsupervised Leiden clustering (resolution 0.5) on single-cell mean intensity of biopsies and control tissues and cell lines, with annotations on right. Lum = luminal, Mes = mesenchymal, Fibro = fibroblast. Colored row labels indicate which biopsy was most dominant for each cluster: Bx2 (orange), Bx3 (blue), or Bx4 (pink). Cluster 16 evenly split between Bx3 and Bx4. (C) Single-cell mean intensity distributions of ER and PCNA staining of cells at 0-25, 25-50, and 50-75 μm away from positive collagen staining. Asterisks indicate significant (p < 0.001) differences in mean intensity between distances (ANOVA). (D) Two views of reconstructed 3D FIB-SEM data from Bx1 showing the intimate relationship between the cancer cells (red and pink), stromal cells (blue and turquoise), and collagen (green). The full volume view on the left shows nanoscale cell-cell interactions of stromal cells surrounding a tumor nest (collagen was not rendered in this image), while the close up view on the right shows a fibroblast-like cell interposed between the tumor and collagen. Scale bars, 5 μm. See also Figure S5 and Table S5.

**Figure 6.**
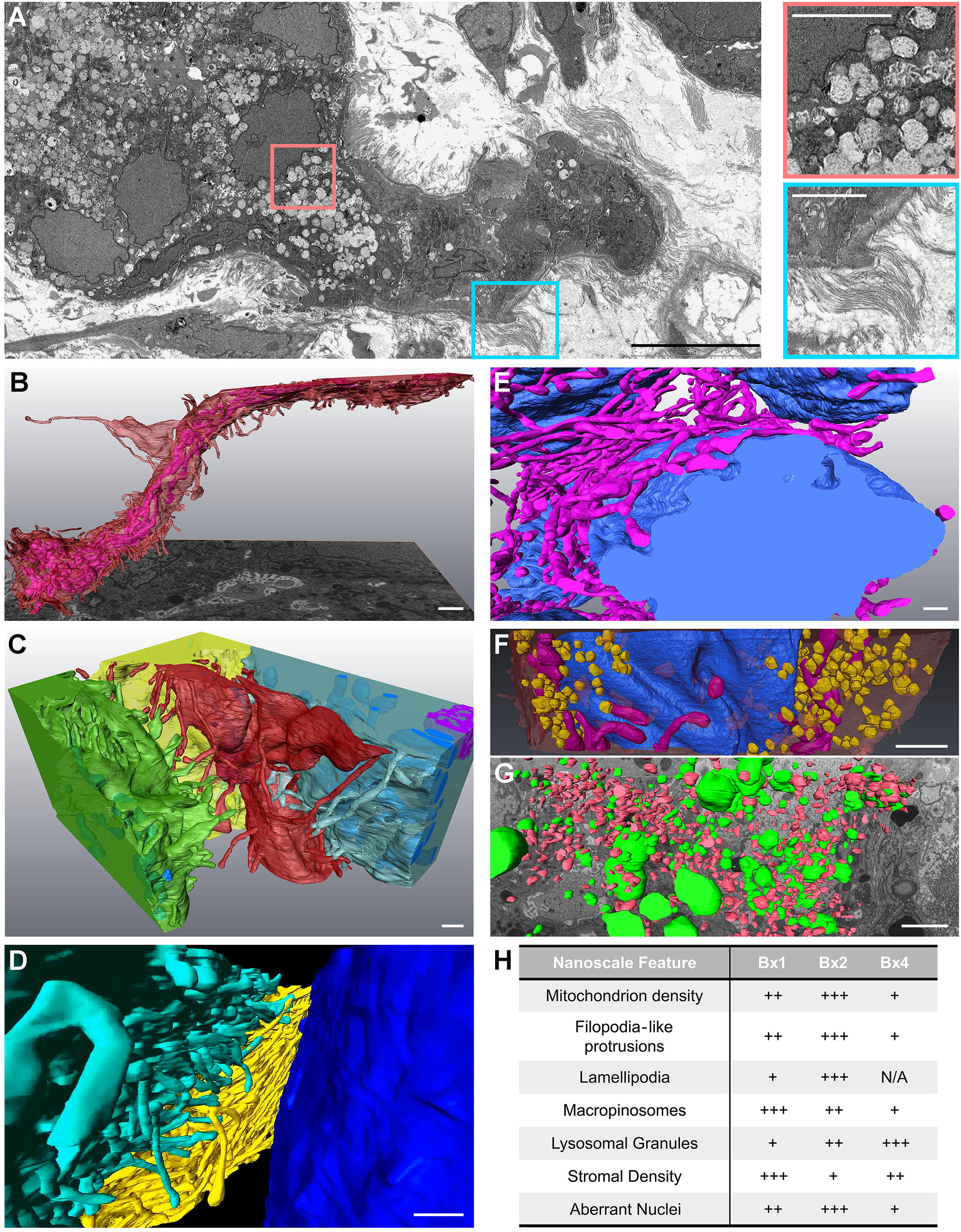
Inter- and intracellular compositions and interactions revealed using focused ion beam-scanning electron microscopy (FIB-SEM) (A) 2D SEM image from Bx4 showing the relationship between tumor cell nests and stromal collagen, along with a high density of extracted lysosomes. Scale bar, 10 μm. The selected insets show these features at high magnification. Scale bars, 3 μm. (B) A side-view of an elongated tumor cell from 3D FIB-SEM of Bx2 showing filopodia-like protrusions (red) and alignment of the internal mitochondria (fuchsia). Scale bar, 1 μm. (C) Additional cells from Bx2 (same red cell as in B) showing paddle-shaped lamellipodia (green cell) and long filopodia-like protrusions (red and blue cells) extending into the stroma and interacting with neighboring cells. Scale bar, 500 nm. (D) Reconstructed 3D FIB-SEM data from Bx1 shows filopodia-like protrusions selectively extending toward neighboring cells and extracellular debris. Scale bar, 1 μm. (E & F) Additional detail from Bx2 (E) and Bx1 (F) of the nuclear invaginations (blue) showing the organization of mitochondria (fuchsia) and macropinosomes (yellow) with respect to nuclear folds. Scale bars, 1 μm. (G) 3D FIB-SEM volume of Bx2 showing large electron-dense lysosomal granules (green) found dispersed between macropinosomes (red). Scale bar, 900 nm. (H) Qualitative summary of ultrastructural feature prevalence within each biopsy. Bx4 scoring of lamellipodia not available. See also Figure S6 and Movies S1-4.

### Genomic evolution between biopsied lesions was substantial

We assessed the genomic landscape, heterogeneity, and evolutionary trajectory of the patient’s cancer by applying targeted DNA and WES to the primary tumor and all four metastatic biopsies. LP-WGS was additionally performed on Bx4 to improve the genome-wide copy number profile. A common set of copy number abnormalities was present in the primary tumor and four biopsies, including amplification of the region on chromosome 11 encompassing the CDK4/6 regulatory partner cyclin D1 (*CCND1*; Figure S3B). Likewise, a subset of somatic mutations identified by WES, including single nucleotide variants (SNVs) and insertion-deletions (Indels), were present in all five tissue samples (Ubiquitous; Figures 3A, S3A). These results indicate that each of the four biopsied lesions arose from a common ancestral cell rather than independent primary tumors. We also identified somatic mutations shared by only a subset of the tissue samples (Shared) as well as somatic mutations unique to each lesion (Private). This intermetastatic heterogeneity between biopsies included differences in biologically-relevant, clinically reported mutations. For example, Bx2 contained a helical-domain hotspot *PIK3CA* mutation (p.E542K; NM_006218:c.1624G>A) that was absent from the primary tumor, Bx1, Bx3, and Bx4. It is noteworthy that Bx2 was acquired during treatment with the combination of fulvestrant, everolimus, and palbociclib, which targets aspects of PI3K signaling. In addition, Bx3 and Bx4 each contained regions of genomic amplification on chromosome 18 (18p11.32) encoding thymidylate synthetase (*TYMS*) and the SRC family tyrosine kinase *YES1* that were not detected in prior biopsies or the primary tumor (Figures 3B, S3B). Analysis of WES revealed the Bx3 amplification was comprised of 8 copies of a 2.3 Mb region of chromosome 18 while the Bx4 amplification was comprised of 14 copies of a 0.7 Mb region. These amplification events were accompanied by increased RNA expression of *TYMS* relative to Bx2 (6.8x for Bx3, 7.2x for Bx4) but less so for *YES1* (2.0x for Bx3, 4.0x for Bx4), supporting a functional consequence of the copy number increase. Importantly, Bx3 and Bx4 were acquired after treatment with capecitabine, which inhibits TYMS.

We used the mutations identified from WES to construct a phylogenetic tree in order to understand the pattern of clonal evolution that gave rise to the intermetastatic heterogeneity between the four biopsied lesions (Figure 3B). This analysis revealed a pattern of branching clonal evolution in which Bx3 diverged from the primary tumor at an earlier evolutionary stage than Bx1, Bx2, and Bx4, which all shared a similar branchpoint. The metastatic lesion sampled in Bx3 was not tracked by CT imaging due to its anatomical location in the spine. However, it first appeared on FDG-PET imaging just one month before the biopsy occurred (Figure S1G), and it was absent from all prior FDG-PET scans (Figures 2, S1A, S1F). These data therefore indicate that the chronological time of emergence of a detectable lesion may not be strictly related to the evolutionary history of the clone. In addition, the phylogenetic relationship between Bx3 and Bx4 implies the amplifications encompassing *TYMS* and *YES1* arose independently in each metastasis.

Genomic differences between the four metastatic biopsies could be due to pre-existing genetic heterogeneity between metastases or to branching clonal evolution during therapeutic treatment. To discriminate between these possibilities, we performed WES on ctDNA collected immediately prior to Bx1 (ctDNA 1) and 23 days after Bx2 (ctDNA 2). At the first ctDNA timepoint, we detected mutations previously identified as private to Bx2, Bx3, or Bx4 (Figure S3A). Furthermore, at the second ctDNA timepoint, we detected mutations previously identified as private to each of the four metastatic biopsies. These results indicate that at least some of the genomic features detected in later biopsies were present prior to initiation of treatment.

### ctDNA increases coincided with both tumor progression and radiation therapy

Treatment response and disease progression was also assessed over the first 32 months of treatment by monitoring ctDNA levels from serial blood samples. Plasma from peripheral blood was subjected to DIDA-Seq for a panel of 53 single-nucleotide variants (SNVs) that were present in the patient’s primary tumor and first two metastatic biopsies.^10^ The average variant allele frequency (VAF) of the panel of SNVs remained consistently below 0.3% of the total cell free DNA in the blood during the monitoring period, with the exception of two transient increases (Figure S2E). The first occurred immediately prior to Bx2, coincident with rising CA 15-3 and CA 27-29 and increasing radiographic size of several metastases, including the Bx2 liver lesion (Figures 2B, 2C, S2E). The increase in ctDNA VAF was greatest for the mutations that were common to the primary tumor and first two biopsies (Bx1_Bx2_Primary, 30% VAF) compared to those private to the metastases (Bx1_Bx2, 3.1%; Bx1, 0.05%; Bx2, 1.3%; Figure S2E). We hypothesize that this variation in VAFs reflects mutational heterogeneity among the diverse metastatic lesions (Figure 2C). The second ctDNA level increase occurred after the patient began a course of palliative radiation therapy to spinal lesions at C2-C5. Interestingly, the VAFs of all SNV groups in the panel increased at this time, including those private to liver lesions Bx1 and Bx2. One possible explanation for this observation is an immune-mediated abscopal radiation effect on lesions both inside and outside the irradiated field.^20^

### Evolving signaling and pathway activities revealed by transcriptional and proteomic analyses

We explored how this patient’s disease evolved over time by applying RNAseq to all four metastatic biopsies. We used the PAM50 subtype gene signature to classify samples into intrinsic molecular subtypes (Parker et al., 2009). The liver biopsies Bx1, Bx2, and Bx4 were all classified as Luminal A, while the bone biopsy Bx3 was classified as Luminal B (Figure S3C). We gained additional context for transcriptional differences between these metastases by comparing RNA transcript levels and pathway activity estimates for the four biopsy samples to other breast cancers from the TCGA-BRCA cohort (Table S2).^9^

We looked for enriched MSigDB Cancer Hallmarks by transcriptomic Gene Set Variation Analysis (GSVA) relative to the TCGA-BRCA Luminal samples.^21,22^ Proliferation, Immune, and Signaling were the most variable MSigDB Hallmark Process categories across the biopsies (Figure 3C). For example, Bx1 exhibited strong enrichment of the transcriptional gene sets “MYC Targets” (Proliferation), “mTORC1” (Signaling), and “PI3K/AKT/mTOR” (Signaling), and reduced “Inflammatory” (Immune) enrichment. Multiple Proliferation and Signaling gene sets decreased in Bx2 relative to Bx1, including the previously mentioned gene sets “MYC Targets”, “mTORC1”, and “PI3K/AKT/mTOR”, as well as “E2F Targets” and late cell cycle gene sets “G2M Checkpoint” and “Mitotic Spindle”. Notably, the Bx2 lesion harboring the *PIK3CA* p.E542K mutation still showed an elevated “PI3K/AKT/mTOR” gene set compared to the TCGA-BRCA samples. Additional gene sets that increased in Bx2 relative to Bx1 include “KRAS Signaling Up” and the Immune gene sets “Interferon-γ”, “IL6/JAK/STAT3”, “IL2/STAT5”, and “Inflammatory”. Compared to the other biopsies, Bx3 exhibited increased enrichment of the Proliferation gene sets “Mitotic Spindle”, “G2M Checkpoint”, and “E2F Targets”, but these sets were not elevated relative to the TCGA samples. In general, Bx4 followed a similar enrichment pattern as Bx1, with elevated “mTORC1” and “MYC Targets” enrichment and reduced Immune gene sets representing interferon, interleukin, and inflammatory signaling.

We also measured total and phosphoprotein levels in Bx1, Bx2, and Bx4 by RPPA (Figure 3E) and assessed pathway activity relative to the TCGA-BRCA cohort using proteomic pathway signatures.^12^ We observed notable variations in pathways representing cell cycle regulation, hormone receptors, and PI3K-AKT signaling between biopsies (Figure 3D). For example, the “G0-G1”, “G2-M”, and “DNA Damage Checkpoint” pathways were increased in Bx1 relative to Bx2 and Bx4, as was phosphorylation of ATM, ATR, CDK1, and WEE1 (Figure 3E), suggesting that Bx1 had activated cell cycle checkpoints in response to DNA damage. In contrast, Bx2 had decreased activity in “G2-M” and increased “G1-S” compared to the other biopsies, consistent with arrest in early cell cycle phases due to treatment with the CDK4/6 inhibitor palbociclib (Figure 2A). Bx4 largely returned to a cell cycle state between that of Bx1 and Bx2, with intermediate activation of “G1-S”, “G2-M”, “Cell Cycle Progression”, and “DNA Damage Checkpoint” pathways.

Aspects of hormone signaling also varied across the biopsies. Consistent with clinical IHC results (Table S1), ER protein levels as measured by RPPA were high in all three biopsies. Interestingly, ER, GATA3, and AR levels all increased in Bx2 compared to Bx1 after treatment with fulvestrant in Phase 1 (Figure 3F). We also observed corresponding increases in “Hormone Signaling” and “Hormone Receptor” protein pathways (Figure 3D) but minimal changes in the GSVA RNA Hallmarks “Estrogen Early” and “Estrogen Late” (Figure 3C), an intriguing finding given that each of these proteins are hormone-regulated transcription factors. Bx4, collected after Phase 4 treatment without hormone suppression, showed continued elevation of the “Hormone Receptor” pathway as well as ER and AR protein levels relative to Bx1. However, GATA3 protein levels, the “Hormone Signaling” protein pathway, and the “Estrogen Early” and “Estrogen Late” GSVA RNA Hallmarks were all downregulated at this final timepoint.

Longitudinal differences in PI3K/AKT/MTOR pathway signaling were also assessed within the proteomic data. While Bx2 was collected after treatment with the mTORC1 inhibitor everolimus and contained the hotspot mutation *PIK3CA* p.E542K, signaling through the “PI3K-AKT” and “TSC-MTOR” protein pathways differed only modestly from Bx1 and Bx4 (Figure 3D). Interestingly, individual protein levels within these pathways did vary substantially across the three biopsies but with a net effect of maintaining similar levels of PI3K and MTOR signaling. It is possible this was a result of feedback loops and compensatory signaling that countered everolimus treatment, as summarized in the Discussion. For example, Bx2 showed decreased mTORC1 complex activity based on decreased S6 phosphorylation at both S235/236 and S240/244 (0.7x and 0.1x vs. Bx1, respectively) but at the same time had evidence of PI3K pathway activation downstream of mTORC2, including increased phosphorylation of AKT (S473: 2.7x vs. Bx1) and its substrates GSK3A/B (S21/S9: 1.7x vs. Bx1), TSC2 (T1462: 1.4x vs. Bx1), and MDM2 (S166: 1.8x vs. Bx1; Figure 3E). Bx4 also had evidence of continued mTORC2 activation, including increased phosphorylation of AKT at S473 (S473: 2.7x vs. Bx1, 1.0x vs. Bx2) and NDRG1 (T346: 1.8x vs. Bx1, 1.6x vs. Bx2), but without an accompanying increase in AKT or mTORC1 substrate phosphorylation, suggesting that mTORC2 was driving PI3K pathway-independent signaling programs in this biopsy.

While limited biopsy tissue precluded generation of RPPA on Bx3, we also used the Intracellular Signaling Protein Panel on Bx2, Bx3, and Bx4 to profile 23 phospho-and total proteins commonly dysregulated in cancer-associated signaling pathways (Figure 3F).^13^ Consistent with RPPA data markers related to the PI3K/AKT/MTOR pathway, the Intracellular Signaling Protein Panel showed Bx2 had the highest phosphorylation of AKT and lowest phosphorylation of S6 relative to the other biopsies. In Bx3, several members of the MAPK pathway were elevated and increased, including p-ERK, p-cRAF, and p-MEK.

Additional insights into tumor evolution can also be gained from integrative analyses that combine multiple datasets from the OMS Atlas. A transcriptional regulator analysis using a molecular interactions network derived from the Pathway Commons resource was used to infer regulator protein activity from the gene expression data.^23^ In addition, integrative analysis of the longitudinal changes in proteomics, phosphoproteomics, gene expression, and transcriptional regulator scores between Bx1 and Bx2 was performed using CausalPath (Figure S3D).^24^ The resulting analysis highlighted dynamic changes in tumor biology between biopsies, including observed changes in PI3K/AKT/MTOR signaling, STAT3/MUC1 signaling, and cell cycle progression. Several changes within the PI3K/AKT/MTOR pathway indicated strong inhibition of MTOR regulator activity (5.1x vs. Bx1) and suggested a possible feedback activation of AKT signaling via mTORC2/Rictor, PDGFR, ER, or ERBB3 (HER3). Increased activity of multiple JAK-STAT family proteins was also observed in the integrative analysis, including JAK2 (1.8x vs. Bx1), phospho-STAT3 (Y705, 1.6x vs. Bx1), and STAT5 (3.2x vs. Bx1), which, together with the oncoprotein mucin 1 (MUC1; Protein 27.1x vs. Bx1; Regulator +3.15 vs. Bx1) constitute a known feed forward loop whereby MUC1 binds to STAT3 to facilitate JAK1 mediated STAT3 phosphorylation.^25^ These observations are consistent with the elevation in “IL6/JAK/STAT3” and “IL2/STAT5” signaling reported by GSVA (Figure 3C). Finally, consistent with decreased GSVA enrichment of “MYC targets” and “E2F1 targets” in the RNAseq, integrative analysis also highlighted a decrease in MYC and E2F regulator activity and E2F1 total protein. Although these observations accompanied by decreased expression of multiple genes involved with cell cycle progression (CCNB1, CDK4, CDK1, CCNE2, CCND3, PLK1) indicates a decline in cell cycle progression, the sharp decrease in cell cycle inhibitors (CDKN1A, CDKN1B, CDKN2A) and lack of changes in RB1 phosphorylation suggest continued proliferative capacity.

### Immune monitoring using mIHC illustrates barriers to T cell activation and tumor immune microenvironment evolution

We utilized a mIHC platform to evaluate the composition and functionality of lymphoid and myeloid lineage immune cells in the primary tumor and biopsies from liver (Bx1, Bx2, and Bx4) and bone (Bx3) (Figure 4). We applied 23-36 antibodies (Figures S4A-E, Table S2) reporting on immune cell identities and functions in single formalin-fixed paraffin-embedded tissue (FFPE) tissue sections and interpreted the results using a computational analysis workflow, as previously described.^14,15^ Changes in immune contexture are noted below, with emphasis on the liver biopsies Bx1, Bx2, and Bx4. We caution that changes in Bx3 relative to other metastases may arise because of its bone origin.

Total immune cell infiltration, as indicated by the percentage of CD45^+^ cells, was similar between the primary tumor and the three liver biopsies, with lowest infiltration detected in Bx2 (Figures 4A, 4B). Analyses of major leukocyte lineages revealed that myelomonocytic cells (macrophages and monocytes) comprised the dominant lineage subgroup in the primary tumor as well as liver Bx1 and Bx2 (Figure 4C) and were notably reduced in Bx4. Deeper analysis of the myeloid lineage using an expanded panel of antibodies revealed that Bx1 contained an increased fraction of immature dendritic cells relative to Bx2 (Figure 4D, blue bar), whereas Bx2 contained increased proportions of macrophages and monocytes (both CD163 positive and negative), with the largest increase in CD163^+^ macrophages in Bx2 (Figure 4D, purple bar). CD163-positivity has been associated with differentiation of myelomonocytic cells towards an alternatively-activated or “M2” type state, an event that is considered to be pro-tumorigenic within solid tumors.^26,27^ CD163 expression on monocytes and macrophages is induced by IL-10 and glucocorticoids and repressed by lipopolysaccharides, TNFα, and IFNγ and is concordant with the observed upregulation in Bx2 of interleukin-containing GSVA gene sets (Figure 3C, Table S2).^28^

The dominance of macrophages and monocytes and relative lack of T cells in the primary tumor, Bx1, and Bx2 was in stark contrast to Bx4, in which there were many more T cells than macrophages and monocytes (Figures 4C, 4E, orange pie slice). This observation is consistent with a mechanism of hepatic siphoning of T cells, in which monocyte-derived macrophages induce Fas-L/Fas mediated apoptosis in CD8^+^ T cells residing within liver metastases.^29^ While we do not have the ability to assess expression of Fas-L on macrophages within the current study, the reciprocity of macrophages and monocytes with T cells in Bx1, Bx2, and Bx4 may be explained by such a mechanism.

We assessed T cell subsets and functionality to gain a deeper understanding of this aspect of immune surveillance. Our analyses showed that only a small fraction of CD3^+^CD4^+^ and CD3^+^CD8^+^ T cells in either the primary tumor, Bx1, or Bx4 expressed the programmed cell death-1 (PD-1) protein that is typically expressed on activated T cells following T cell priming or persistent antigen exposure (Figure 4F).^30^ Low expression of PD-1 by T cells at these timepoints is consistent with impaired T cell-mediated immune response. However, the T cell status was markedly altered in Bx2 (Figures 4C, S4F). Notably, while T cells were least abundant in Bx2 compared to Bx1 and Bx4, the largest fraction of CD3^+^CD4^+^ and CD3^+^CD8^+^ T cells expressing PD-1 was observed in Bx2 (Figure 4F), coincident with a relatively reduced presence of FoxP3^+^CD4^+^ Tregs and an expanded population of Th17 CD4^+^ T cells (Figures 4G, 4H).

We measured PD-1 and eomesodermin (EOMES) expression to further audit CD8^+^ T cell differentiation and functional status. These analyses revealed that the primary tumor contained predominately PD-1^-^EOMES^-^ and PD-1^+^EOMES^-^ CD8^+^ T cells, likely reflecting naïve and early effector subsets, respectively. Evolution of CD8^+^ T cells in liver Bx1, Bx2, and Bx4 indicated progressive loss of PD-1^-^EOMES^+^ (late effector, green) and PD-1^+^EOMES^+^ (exhausted, red) subsets, with replacement by likely naïve (purple) PD-1^-^EOMES^-^ CD8^+^ T cells (Figure 4G).

The Bx3 bone metastasis differed from the primary tumor and liver metastases, with the bone having the highest percentage of CD45^+^ leukocytes (Figures 4A, 4B), with comparatively high percentages of granulocytes, dendritic cells, and CD20^+^ B cells (Figure 4C). However, like liver Bx4, bone Bx3 contained a prominent granulocyte infiltrate that most likely are predominantly neutrophils. Neutrophils can exert significant pro-metastatic activities, including suppressive effects on T cells, and are associated with poor prognosis in many solid tumors, including breast cancer.^31–35^

### Tumor and stromal interactions defined using CycIF and FIB-SEM

We used a CycIF analysis platform with probes for 24 proteins (Table S2) as previously described to assess the tumor and stromal composition and organization of all four biopsied metastatic lesions.^17,18^ Probes to collagen I and collagen IV were included to enable assessment of the extracellular matrix (ECM). All four biopsies were analyzed along with control biospecimens prepared from normal breast tissue, tonsil, and six cell lines representing basal-like (HCC1143, HCC3153), claudin-low (MDAMB436), luminal (T47D), and HER2 positive (BT474, AU565) breast cancers in order to enable subpopulation analysis within the context of known biology for this luminal breast cancer (Figure S5B-E). Figure 5A shows illustrative images for Bx2, Bx3, and Bx4. Bx1 was analyzed before probes for the collagens were available and so is not shown. The measurements of cell protein expression levels for each segmented cell in the four biopsies and associated control samples were organized into 17 clusters, as described previously (Figures 5B, S5A). Three of the stromal clusters (clusters 11, 13, and 14) and five of the tumor clusters (clusters 0, 1, 5, 7, and 9) comprised major subpopulations in Bx2, Bx3, and Bx4. The three stromal clusters were identified as fibroblast-like cells that differed in levels of vimentin (VIM). The markers for endothelial cells (CD31) and macrophages (CD68) were excluded from cluster analysis due to loss during staining of the normal breast and tonsil tissues, normally used to control these markers during normalization. We confirmed the presence of these populations using manual gating (Figure S5F). The tumor clusters expressed CK7/CK19 but expressed different levels of ER, EGFR, and CK8 (Figure 5B). An additional proliferative cluster (cluster 16) appeared in Bx3 and Bx4 that was comprised of tumor and stromal cells expressing high levels of Ki67 and/or PCNA. Tumor, endothelial (CD31+), immune (CD45+), and fibroblast (VIM+) cells are indicated in Figure 5A as color coded cell segments.

Spatial analyses indicated that the tumor cells were formed into nests surrounded by immune, fibroblast, and endothelial cells as well as collagen I and collagen IV deposits. This was observed in all biopsies but was pronounced in Bx3 from the bone. Quantitative analysis of the expression of ER and PNCA in the tumor nuclei in Bx2, Bx3, and Bx4 as a function of distance to the collagen I rich tumor nest boundaries showed that the cells expressing higher levels of ER and PCNA were closest to the collagen I rich stromal boundary (Figure 5C) and other stromal compositions (Figure S5G). This is consistent with previously reported increases in estrogen-mediated proliferation by interactions with stromal cells and collagen I.^36,37^

We explored the features of the tumor-tumor and tumor-stromal interactions in Bx1, Bx2, and Bx4 at ∼4 nm resolution using FIB-SEM, as previously described.^19^ Insufficient material was available from Bx3 to allow SEM analysis. Computational renderings of the 3D images of a segment of Bx1 (Movies S1, S2) and a segment of Bx2 (Movie S3) reveal details of interactions between tumor and stromal cells and ECM proteins that cannot be seen using light microscopy. For example, Movie S2 and selected views in Figure 5D show a previously unappreciated lattice-like structure for the fibroblast-like cells surrounding tumor cell clusters and an intricate interaction pattern between these cells, collagen bundles, and the tumor cells on the nest boundaries. The production of collagen by tumor associated fibroblast-like cells is particularly apparent in the 2D SEM image of Bx4 (Figure 6A). Interestingly, the 3D images suggest that the fibroblast-like cells are interposed between the tumor cells and the collagen bundles in most cases, raising the issue of how collagen stiffening leads to more aggressive tumor behavior.^38^

These images also show the robust manifestation of ∼100 nm diameter, micrometers long filopodia-like protrusions (FLPs) and lamellipodia that appear to establish interactions between adjacent tumor cells and between the tumor and the stromal microenvironment (Figures 6B-D, Movie S3). Published work and our own studies in model systems show that these protrusions have actin-rich cores that are decorated with receptor tyrosine kinases that are transported along the FLPs by the actin-motor protein Myosin-X.^39^ Dynamic *in vitro* images acquired using stochastic optical reconstruction microscopy (STORM) show that cultured, FLP-rich SKBR3 breast cancer cells respond to cytokines like epidermal growth factor (EGF) by rapidly decreasing protrusion length in ways that cause cells to move toward the anchored ends of the protrusions (Figure S6B, Movie S4). This indicates the possibility of protrusion-mediated signaling whereby the long and dynamic FLPs mediate both proximal and distal interactions and directed movement. This mechanism might provide the force needed to produce the elongated tumor cell shown in Figure 6B with mitochondria aligned along its long axis and inserted into nuclear folds (Figure 6E). Published studies also suggest the possibility that FLPs mediate protein transport between cells.^40^ Figure 6C and Movie S3 also show evidence of lamellipodia surrounding a region of apparent cell debris, suggestive of the structures observed in model systems that enable nutrient scavenging via macropinocytosis.^41,42^

### Intracellular nanobiology defined by FIB-SEM

The 3D FIB-SEM images of cancer cells in Movies S1-3 also provide important information about intracellular structures and interactions that may influence cell function and therapeutic response. For example, Movie S3 and Figures 6B and 6E from Bx2 show mitochondria aligned along the length of an elongated cell and insinuated into nuclear folds, the latter increasing the potential for nuclear-mitochondria signaling. The resulting increased mitochondria-nucleus proximity might alter aspects of DNA damage repair and/or reactive oxygen species (ROS) signaling.^43,44^

Movies S1 and S3 as well as Figures 6C, 6F, and 6G depict a high abundance of lamellipodia and macropinosomes, implicating nutrient scavenging via macropinocytosis as a possible tumor survival mechanism.^45,46^ Macropinocytosis is a non-selective endocytosis process that enables the uptake of nutrients and proteins from the intercellular space, including those from nearby dying cells.^45^ Uptake is mediated by actin-rich lamellipodia that engulf and internalize extracellular materials. This process has been implicated as a survival mechanism in amino acid-poor environments.^47^ Qualitative analyses of 2D SEM images show that the frequency of macropinosomes decreases progressively from Bx1 to Bx4.

Movie S1 from Bx1, Movie S3 and Figure 6G from Bx2, and Figure 6A from Bx4 show a high prevalence of densely stained vesicles that appear to be lysosomes, in contrast to the smaller electron-lucid macropinosomes. These acidic vesicles are known to accumulate weak bases, including some cancer drugs, via a process called lysosomotropism. In this process, basic drugs become protonated and trapped within the acidic interior of lysosomes.^48^ Analysis of 2D SEM images show that the density of these vesicles increases progressively from Bx1 to Bx4 (Figures 6H, S6A).

Figure 6H presents a qualitative summary of the prominence of the nanoscale features described above in Bx1 and Bx2 made by visual analysis of large format 2D SEM images (Figure S6A) and informed by 3D FIB-SEM images of selected features as described above.

## Discussion

This report describes our efforts to comprehensively catalog the cellular, molecular, and organizational composition of four tumor biopsies collected over a 3.5-year period from a single patient with metastatic breast cancer. The larger goal of this study and the SMMART and HTAN programs is to improve treatments for metastatic cancers in individual patients by identifying and opportunistically acting on therapeutic vulnerabilities and mechanisms of resistance as they present before and during extended treatments. A truly personalized approach such as this requires advanced methods for collecting and analyzing both clinical and molecular data, coupled with the ability to interpret results in the context of longitudinal samples from the same patient. With the generation of this OMS Atlas, we have now demonstrated the feasibility of the biospecimen acquisition, management, and analysis necessary to achieve this goal.

As a proof-of-principle, we discuss below insights derived from analysis of the OMS Atlas into mechanisms of drug resistance and response that arose in biopsies taken over the course of four phases of treatment. However, we acknowledge the difficulty in assigning specific responses of the tumor to individual drugs within a multi-drug treatment regimen, especially for drug combinations targeting overlapping biological pathways. Moreover, working with a single human subject precluded the implementation of hypothesis testing that is *de rigueur* in experimental cell lines and animal models and would more definitively support our proposed mechanisms. These factors demonstrate the challenges of implementing this type of program and analyzing N- of-1 data in a real-world clinical setting. Nonetheless, we were able to identify several potential resistance mechanisms and new therapeutic vulnerabilities in each biopsy by combining our broad array of analytics with curated public datasets and published literature (Figure 7). It is important to note that the research data were not used to guide treatment and are instead presented here *post hoc*.

**Figure 7.**
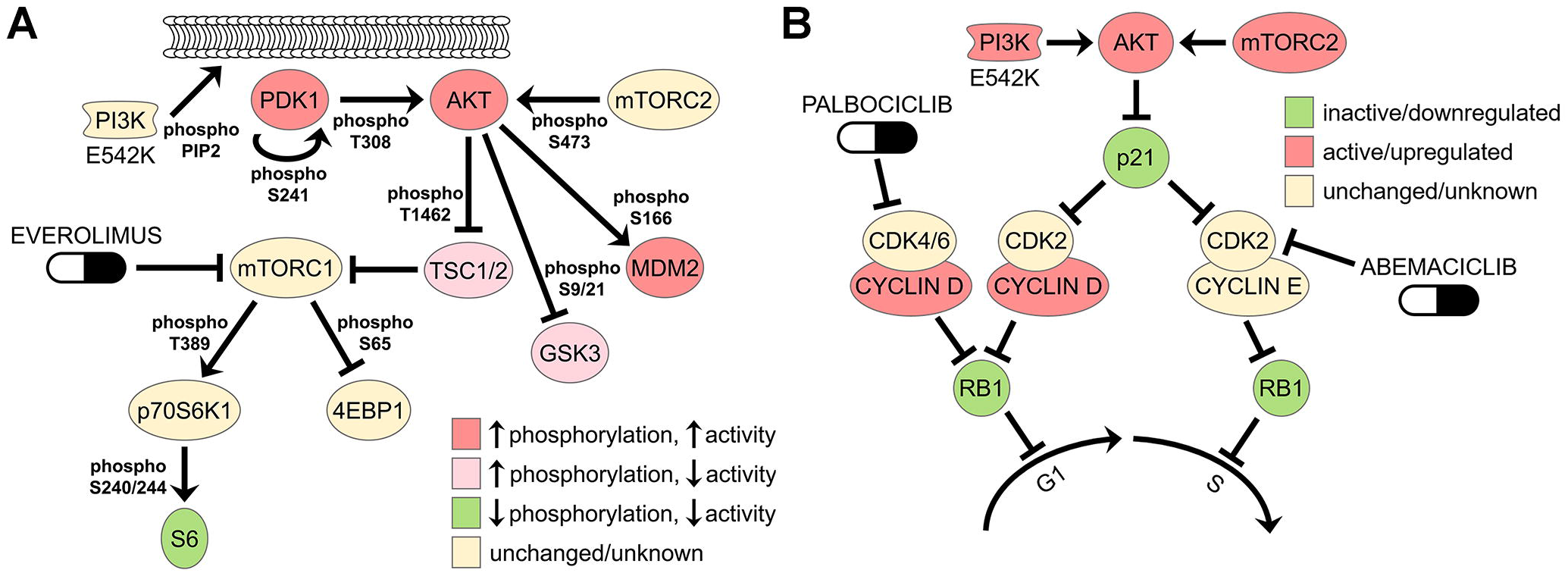
Mechanisms of therapeutic resistance and response suggested by analyses of RPPA data (A) Phosphorylation and inferred activation of the PI3K-AKT-mTOR pathway effected by everolimus in Bx2. Decreased phosphorylation of S6 downstream of mTORC1 likely resulted from everolimus inhibition, but increased phosphorylation of proteins downstream of PI3K and AKT, possibly through mutant PI3K E542K activity and/or feedback signaling to mTORC2, may have provided continued oncogenic signaling in the presence of this drug. Proteins are noted as increased activating phosphorylation (> 1.4x; red), increased inhibitory phosphorylation (>1.4x; pink), decreased activating phosphorylation (< 0.7x; green), unchanged phosphorylation (yellow), or unknown phosphorylation status (white). Changes in phosphorylation in Bx2 vs. Bx1: PDK1 = 1.45x; AKT T308 = 1.20x; AKT S473 = 2.69x; TSC2 = 1.43x; GSK3A/B = 1.71x; MDM2 = 1.75x; p70S6K = 0.92x; 4EBP1 = 1.37x; S6 S235/236 = 0.69x; S240/244 = 0.14x. (B) Activation status for cell cycle regulatory pathways effected by palbociclib in Bx2, as inferred by total and phospho-protein levels. Palbociclib blocks cell division in responsive cells by inhibiting CDK4/6 phosphorylation of RB1, but Bx2 had continued high levels of phospho-RB1 and cell proliferation under treatment with this drug (RB1 P-S807/811 = 0.98x vs. pre-treatment Bx1). This is possibly due to degradation of the CDK2 inhibitor p21 (0.50x vs. Bx1) by activated PI3K/AKT signaling (see panel A), which would activate canonical cyclin E/CDK2 complexes to drive cells through G1-S. Alternatively, cell division might be proceeding through the formation of non-canonical cyclin D1/CDK2 complexes due to amplified *CCND1* (Figure S3B), high levels of cyclin D1 protein (1.67x vs. Bx1), and low p21. CDK2 activation can be countered with the broad-spectrum CDK inhibitor abemaciclib. Inferred activation status is based on total protein levels or phosphorylation and is designated as relative increases (red), decreases (green), or unchanged/unknown (yellow). See also Figure 3E and Table S2.

Phase 1 treatment consisted of a combination of fulvestrant, palbociclib, and everolimus and is supported by the findings in Bx1 of high ER protein expression, wild type *ESR1*, and two intact copies of wild type *RB1*. Bx2 was taken after CT imaging and blood biomarkers demonstrated that this combination was no longer adequately controlling disease in a few lesions after about 8 months (Figure 2). Analyses of Bx2 taken from one progressing lesion revealed multiple possible mechanisms for resistance to this combination. For example, one well-known mechanism by which cells become resistant to everolimus and other mTORC1 inhibitors is through mTORC2 activation.^49,50^ Consistent with this, phospho-protein analyses revealed that S6 phosphorylation sites in Bx2 were decreased, which is evidence for continued inhibition of mTORC1 by everolimus. At the same time, Bx2 had increased phosphorylation at an mTORC2 site on AKT (S473) and of multiple AKT substrates that together are predicted to maintain oncogenic PI3K/mTOR signaling in the presence of this drug (Figure 7A). Everolimus efficacy might also have been reduced by a *PIK3CA* p.E542K activating mutation unique to Bx2, which is known to strongly activate the PI3K/AKT/MTOR pathway.^51^ Indeed, this variant was among the SNVs monitored in serial blood samples by DIDA-Seq (Figure S2E) and was only detected in the ctDNA significantly above background after seven months on Phase 1 therapy (0.06% VAF, p=0.0071, Weitzman overlapping coefficient), indicating that this mutation may have emerged due to selective pressure from one or more Phase 1 drugs.

Our analyses also yielded evidence suggesting mechanisms of resistance to the CDK4/6 inhibitor palbociclib in Bx2. An important biomarker for palbociclib efficacy is phosphorylation of RB1, which at high levels promotes cell cycle progression by activating E2F and when low results in RB1-mediated cell cycle arrest.^52^ Palbociclib blocks cell division by inhibiting the G1 cyclin-dependent kinases CDK4 and CDK6, preventing them from phosphorylating and inhibiting RB1. Bx2 had a level of RB1 phosphorylation comparable to that of the pre-treatment Bx1 (P-S807/S811 was 1.0x vs. Bx1), indicating that a loss of palbociclib efficacy may have contributed to tumor progression at the end of Phase 1. Loss of *RB1* has been shown to be an important driver of CDK4/6 inhibitor resistance in multiple clinical trials,^53^ but this gene was intact and unmutated in Bx2. Instead, protein profiling of key cell cycle regulators revealed that this biopsy might have bypassed palbociclib by activating CDK2, an additional cyclin-dependent kinase that inhibits RB1 but is not inhibited by palbociclib (Figure 7B). For example, protein levels of the CDK2 inhibitor p21 were decreased 2x from Bx1 to Bx2, which may result in increased activity of CDK2/cyclin E complexes that phosphorylate RB1 to drive the cells through G1-S.^54–56^ p21 protein levels might be reduced by activated PI3K/AKT, as this pathway keeps p21 levels low in CDK4/6 inhibitor resistant cells.^56^ Tumor cells can also adapt to palbociclib via non-canonical binding of CDK2 to cyclin D1, which occurs in cells with high cyclin D1 and activated PI3K.^54^ Bx2 had both increased cyclin D1 protein levels (1.7x vs. Bx1) and increased PI3K/AKT signaling activity compared to Bx1, consistent with this resistance mechanism. The CDK4/6 inhibitor abemaciclib has a broader spectrum of activity, including CDK2,^57^ and so might be expected to be effective in cases where palbociclib escape occurs via CDK2 activation. Indeed, abemaciclib administered subsequent to the period covered by this study did show efficacy (data not shown). Finally, sequestration via lysosomotropism has been implicated as a mechanism of resistance to CDK4/6 inhibitors.^58,59^ This mechanism is suggested by the increase in lysosomes from Bx1 to Bx2 revealed by FIB-SEM (Figures 6G, 6H, and S6A; Movies S1 and S3) and so may have contributed to the palbociclib resistance observed in the lesion from which Bx2 was taken. Interestingly, the anti-infective, immunosuppressive agent hydroxychloroquine, sometimes used to counter treatment induced rashes, is reported to have lysosomotropic activity and thus might reduce treatment efficacy when co-administered with any basic drug, including CDK4/6 inhibitors.^60^ Recent studies of lysosomotropism-mediated doxorubicin resistance in angiosarcoma cells suggest that this potential resistance mechanism can be countered by treating with the beta adrenergic receptor (beta-AR) antagonist propranolol, which acts through a beta-AR-independent mechanism to increase cytoplasmic doxorubicin concentrations and decrease lysosomal accumulation.^61^

Capecitabine was administered in Phase 3 along with pembrolizumab, enzalutamide, and fulvestrant. While this combination initially reduced tumor burden, evidence of disease progression was apparent after about 20 months (Fig 2B). Analysis of Bx3 revealed a focal amplification on chromosome 18 containing *TYMS,* an enzyme involved in nucleotide biosynthesis that is inhibited by capecitabine, and concomitant increased levels of *TYMS* gene expression. As overexpression of *TYMS* is a potential resistance mechanism to capecitabine, this amplification may have given Bx3 a relative fitness advantage under the selective pressure of capecitabine treatment.^62^ This would explain the temporally late emergence of a clone that branched off early in the evolutionary process (Figure 3B). *YES1,* located in close genomic proximity to *TYMS,* was also amplified in Bx3 but was not as highly expressed in that biopsy. Amplification of *TYMS* and *YES1* arose independently in Bx4, presumably due to the earlier exposure to capecitabine, but in this biopsy both genes were overexpressed, indicating that *YES1* overexpression may have provided a growth advantage to the lesions from which Bx4 arose months after cessation of capecitabine (Figure 2A). YES1 is a SRC family tyrosine kinase and a target of the broad-spectrum tyrosine kinase inhibitor dasatinib, so inhibition of YES1 may be considered as a possible orthogonal therapeutic strategy for patients who become resistant to capecitabine via amplification of *TYMS/YES1*. However, dasatinib was administered subsequent to the period covered by this study and did not show efficacy (data not shown), arguing against this strategy.

Comparative analyses of the primary tumor and serial biopsies suggested several mechanisms that shaped the immune contexture. The most significant was associated with treatment with the CDK4/6 inhibitor palbociclib at the time of Bx2. Our mIHC analyses showed substantially increased abundance of macrophages/monocytes, increased proportion of Th17 cells, and decreased Tregs in Bx2 compared to Bx1 and Bx4 (Figure 4). Th17 cells and Tregs arise from a common precursor but have opposing functionality upon terminal differentiation, with Th17 cells promoting and Tregs dampening antitumor immunity.^63^ This suggests that the relatively high frequency of Tregs in the primary tumor and Bx1 may have contributed to reduced T cell activation, as detected by lack of PD-1 expression (Figure 4G). Conversely, the Th17 dominance over Tregs in Bx2 may have been involved in supporting T cell activation, as evidenced by increased PD-1 expression. These changes were coincident with substantial increases in IFNγ signaling as well as signaling through numerous interleukins and STATs, as revealed by gene and protein expression profiles (Figure 3C, Table S2). These changes are consistent with studies of model mammary tumor systems that showed that CDK4/6 inhibitors promote antitumor immunity by stimulating type III interferons and suppressing proliferation of Tregs, thereby promoting T cell-mediated clearance of tumor cells.^64^ Our observations relating to Bx2, coupled with the increase of PD-1 expression in T cells, supported the utility of an immune checkpoint inhibitor, which was administered during Phase 3 treatment. However, the immune contexture changed again after discontinuation of treatment with palbociclib, with Bx4 showing a decreased number of macrophages/monocytes, an increased number of T cells, only a small fraction of which expressed PD-1, and a reduced fraction of Th17 T-cells. Although Bx4 also contained reduced percentages of Tregs (similar to Bx2) and the highest proportions of Th1 differentiation (Figure 4G), there was low PD-1 expression on T cells (Figure 4F). These results indicate a lack of T cell priming in Bx4, potentially due to low neoantigens and antigen presentation as likely barriers to functional anti-tumor immunity, as opposed to T cell-mediated suppression, which could have been underlying the lack of T cell responses observed in the primary tumor and Bx1.

Analyses using CycIF and FIB-SEM showed tumor cells were organized into nests surrounded by stromal cells and substantial collagen I deposits (Figures 5, 6), suggesting that the lack of neoantigens and/or antigen presentation inferred from immune profiling may be caused, at least in part, by a biophysical barrier that diminishes tumor-immune cell interactions. Active dead cell scavenging via tumor macropinocytosis as suggested by FIB-SEM (Figure 6) may further diminish communication of neoantigens to the immune cells by competing with dendritic cells for exogenous antigens from dying tumor cells.^65^ Interestingly, the FIB-SEM analyses also showed that the tumor-stromal interactions at the edges of tumor nests were remarkable in their complexity, with filamentous fibroblast-like cells interspersed between the tumor cells and collagen depositions. How immune cells interact with these structures remains to be elucidated, as are the mechanisms by which the stromal barriers stimulate the increased expression of ER and PCNA in closely proximal tumor cells (Figure 5D). From a technical perspective, the complex cellular interactions revealed by FIB-SEM illustrate the difficulties of properly segmenting individual cells during multiplex imaging of 2D sections using mIHC or CycIF (Figure S6C) and of dissociating tightly interacting and potentially fragile cells for single cell analyses.

The aspects of macropinocytosis and possibly forced mitochondria-nuclear interactions revealed in 2D and 3D FIB-SEM images of cancer cells (Figure 6, Movies S1-S3) also suggest several therapeutic actions. For example, the reliance of cells on macropinocytosis as illustrated in Figures 6C, 6F, and 6G suggests that treating with protein-conjugated drugs might convert this survival mechanism into a therapeutic vulnerability. We speculate that this mechanism may have been partly responsible for the control achieved by treatment with liposomal doxorubicin during Phase 3. The apparent forcing of mitochondria into nuclear folds by the filopodia-like protrusion mediated movement suggested by Figure 6E may lead to increased DNA damage related crosstalk between the nucleus and mitochondria and increased reactive oxygen signaling.^44,66^ We speculate that this might be countered therapeutically by attacking either reactive oxygen signaling or by inhibiting FLP function.

Overall, this OMS Atlas shows both the promises and challenges of elucidating evolving resistance mechanisms and new therapeutic vulnerabilities in individual patients. Although this type of complex, comprehensive, and integrative analysis provides meaningful insight into mechanisms of tumor response and resistance, we acknowledge that these methods are currently too complex to be widely applied. However, once the utility of each assay platform is established, workflows can be dramatically simplified and turn-around-times shortened. Our work shows that further development of analytical methods to integrate and interpret multi-platform omic and imaging datasets is clearly needed for both the clinical and research communities and we believe that the OMS Atlas can serve as a resource in that effort.

A further challenge of this approach of interrogating evolving resistance mechanisms is the remarkable intra- and intermetastatic heterogeneity that exists in some cancers, which is critical to understand for optimal disease management.^67^ A biopsy of a single metastatic lesion at any single time point is unlikely to provide a comprehensive picture of the entire heterogeneous disease within a patient, nor may it provide sufficient tissue to fully capture the heterogeneity even within the biopsied lesion. This complicates attempts to use the integrative analyses described herein to interpret biological differences between metastases and across a treatment timeline, as any observed changes may due to sampling bias and not functionally important. This is a fundamental limitation of any biopsy-based analytical strategy. The continued advancement of assays that report on overall tumor composition across multiple lesions, such as peripheral blood assays, is one potential avenue toward understanding heterogeneous disease burden. Indeed, our observation that radiation induced a transient increase in ctDNA in peripheral blood suggests that patients undergoing radiotherapy might have circulating tumor nucleic acids and proteins at sufficient quantities for both practical diagnostic measurement and for revealing latent, low-level molecular changes in unbiopsied lesions in near real time.

In conclusion, there are significant challenges to be overcome for the full realization of adaptable, personalized cancer treatments based on the complex tumor and microenvironmental mechanisms of therapeutic response in each patient. Toward that end, the present study shows that omic and image analyses from serial biopsies can be safely implemented and that integration and interpretation of the resulting data provides insight into the diverse resistance and response mechanisms that manifest during an extended course of treatment.

## Acknowledgments

Work at OHSU on this project was carried out with major support to the OHSU SMMART (Serial Measurement of Molecular and Architectural Responses to Therapy) Program from the National Institutes of Health (NIH), National Cancer Institute (NCI) Human Tumor Atlas Network (HTAN) Research Center (U2C CA233280), and the Prospect Creek Foundation. The program was initiated with support from a Stand Up to Cancer-American Association for Cancer Research Dream Team Translational Cancer Research Grant, SU2C-AACR-DT0409. Additional support came from the OHSU Brenden-Colson Center for Pancreatic Care, the W. M. Keck Foundation, the NIH/NCI Cancer Target Discovery and Development (CTD^2^) (U01 CA217842), a NIH/NCI Cancer Systems Biology Consortium Center (U54 CA209988), NIH/NCI U01 CA224012 (to L.M.C.), a SBIR (R44 CA224994) (to K.C.), the Damon Runyon Cancer Research Foundation (to X.N.), and the M. J. Murdock Charitable Trust. Sequencing and multiscale microscopy was supported by the Knight Cancer Institute Cancer Center Support Grant (5 P30 CA69533). Electron microscopy was performed at the Multiscale Microscopy Core; light microscopy was performed using equipment in the Advanced Light Microscopy Core, both OHSU University Shared Resource Cores. Short read sequencing assays were performed by the OHSU Massively Parallel Sequencing Shared Resource.

We acknowledge the following teams for their assistance with this study:

Clinical: Raymond Bergan, Christopher Corless, Alexander R. Guimaraes, Ben Kong, Zahi Mitri, and George Thomas.

Research Operations: Heidi S. Feiler, Joe W. Gray, Brett E. Johnson, Jamie Keck, Taylor Kelley, Marlana Klinger, Annette Kolodzie, Gordon B. Mills, Max Morris, Anastasiya Olson, Swapnil Parmar, Kiara Siex, Jayne M. Stommel, and Leanna Williams.

Information management: Imogen Bentley, Patrick Leyshock, Georgia Mayfield, Souraya Mitri, Damir Sudar, Matt Viehdorfer, and Christina Zheng.

Nucleic acids: Christopher Boniface and Paul T. Spellman.

Proteins: Aurora Blucher, Todd Camp, Marilyne Labrie, and Jinho Lee

Omics analysis: Özgün Babur, Allison L. Creason, Emek Demir, Joseph Estabrook, Jeremy Goecks, Laura M. Heiser, Janice Patterson, and Julia Somers.

Image analysis: Erik Burlingame, Young Hwan Chang, and Guillaume Thibault.

mIHC: Teresa Beechwood, Konjit Betre, Courtney B. Betts, Gina Choe, Lisa M. Coussens, Giovanney Gonzalez, Nell Kirchberger, Lauren Maloney, and Shamilene Sivagnanam.

CycIF: Elmar Bucher, Koei Chin, Zhi Hu, and Jennifer Eng.

FIB-SEM: Steven Adamou, Dylan Blumberg, Cecilia Bueno, Kaylyn Devlin, Yingsi Gao, David Kilburn, Moqing Liu, Kevin Loftis, Jessica L. Riesterer, Hannah Smith, Rebecca Smith, Kevin Stoltz, and Erin S. Stempinski.

STORM: Xiaolin Nan and Jing Wang.

## Author Contributions

Conceptualization, R.B., G.B.M., and J.W.G.; Methodology, B.E.J., J.M.K., S.P., A.K., K.C., X.N., L.M.H., P.T.S., E.D., L.M.C., C.C., J.G., and J.W.G.; Software, A.L.C., P.L., S.S., D.S., G.Thibault, E.D., and Y.H.C.; Formal Analysis, A.L.C., A.B., C.B., E.Bucher, E.Burlingame, J.Eng, J.Estabrook, M.L., J.L., J.P., S.S., J.S., and G.T.; Investigation, B.E.J., J.M.K., B.L.K., S.P., C.B.B., C.B., T.C., K.C., Z.H., J.Eng, J.L.R., X.N., G.Thomas, A.R.G., and C.C.; Resources, R.B. and Z.M.; Data Curation, A.L.C., J.M.K., S.P., S.M., and C.Z.; Writing – Original Draft, B.E.J., A.L.C., J.M.S., J.M.K., S.P., C.B.B., K.C., J.L.R., J.G., and J.W.G. Writing – Review & Editing, B.E.J., A.L.C., J.M.S., J.M.K., C.B.B., A.B., K.C., A.K., B.L.K., M.L., J.L.R., L.M.C., J.G., R.B., Z.M., G.B.M., and J.W.G.; Visualization, A.L.C., J.M.S., S.P., C.B.B., E.Burlingame, K.C., J.Eng, M.L., J.P., J.L.R., and X.N.; Supervision, C.C., J.G., R.B., Z.M., G.B.M., and J.W.G.; Project Administration, B.E.J., S.P., H.S.F., and A.K.; Funding Acquisition, L.M.C., C.C., J.G., R.B., G.B.M., and J.W.G.

## Tables

None

## Methods

Patient Consent and Biospecimen Collection

This study was approved by the Oregon Health & Science University (OHSU) Institutional Review Board (IRB). All biospecimens and data were acquired and analyzed under the OHSU IRB-approved protocols *Molecular Mechanisms of Tumor Evolution and Resistance to Therapy* (IRB#16113) and *Reconstructing the Tumor Genome in Peripheral Blood* (IRB#10163). Participant eligibility was determined by the enrolling physician and informed written consent was obtained prior to all study protocol related procedures.

### Resource Availability

All primary datasets from clinical and exploratory analytics generated during this study are available through the HTAN Data Coordinating Center (https://humantumoratlas.org/) as patient HTA9_1 in the OMS Atlas.

The published article includes all clinical metadata analyzed during this study (Table S1).

This study did not generate new unique reagents.

### Radiology

FDG-PET/CT imaging was performed according to the standard institutional protocol, with patients fasting for 6 hours following 24 hours of rest. Prior to the examination and FDG injection, blood glucose levels were confirmed to be less than 200 mg/dL. The patient received a dose of ^18^F-FDG of 370 to 555 MBq (10-15 mCi) on the basis of body weight. After an uptake period of 90 minutes, a vertex-to-mid-thigh FDG-PET/CT scan was performed using 3 min/bed position on a CTI Biograph duo PET/CT scanner (Siemens Medical Systems, Hoffman Estates, Illinois, USA) or a CTI Biograph TruePoint 40 PET/CT scanner (Siemens Medical Systems, Knoxville, Tennessee, USA). CT imaging was performed according to the standard institutional protocol from clavicles to mid-thigh on a Phillips Brilliance CT 128slice helical scanner (Philips Medical Systems, Amsterdam, NE).

Pre- and on-treatment FDG PET/CT studies were reviewed by an expert nuclear medicine physician with analysis performed by a body imager with 15 years of experience in oncologic imaging. Target lesions were selected by having maximum standard uptake values (SUVmax) greater than normal mediastinum average (lymph nodes), and greater than background liver SUV (liver lesions) and were recorded on the pre- and on-treatment scans at the same tumor region. Image analysis was performed using syngo via advanced visualization software (Siemens Healthcare GmbH, Erlangen, Germany) and Horos visualization software (Horos, Lausanne, Switzerland). All lesions meeting these criteria were recorded both on FDG-PET/CT and combined with long axis measurements (e.g., liver, splenic, lung lesions) and long and short-axis measures (lymph nodes) at all time points during the care of the patient. All SUVmax measures were normalized by subtracting the mean background SUVmax from the organ of origin (e.g., mediastinum or liver).

### GeneTrails^®^ Solid Tumor Panel

The GeneTrails Solid Tumor Panel was performed by the OHSU Knight Diagnostic Laboratories on genomic DNA extracted from macro-dissected, tumor-rich regions of FFPE. Next-generation sequencing libraries were prepared using custom QIASeq chemistry (QIAGEN) with multiplexed PCR and sequenced on an Illumina NextSeq500/550. The DNA library was generated by 9,229 custom-designed primer extension assays covering 613,343 base pairs across 225 cancer-related genes (including whole exons of 199 genes and hotspot regions of 26 genes). This panel is routinely sequenced to an average read depth of >2,000, providing high sensitivity for SNVs, short insertions/deletions, and copy number alterations. Variants were identified using both FreeBayes and MuTect2 algorithms in a custom sequencing analysis pipeline.

### Blood collection and DNA isolation for WES, DIDA-Seq, and LP-WGS

Up to 40 mL (range 6-40 mL) of blood were collected in 5x 6-mL or 4x 10-mL, purple-capped EDTA tubes. Consistent with published recommendations, blood plasma was isolated within 6 h of collection by first spinning whole blood at 1000g for 10 min, separating the top plasma layer into 1 mL aliquots, then spinning those aliquots at 15,000g for 10 min, transferring the supernatant to cryovials, and storing at −80°C.^68^ DNA extraction of tumor tissue from FFPE was carried out using QIAGEN FFPE DNA extraction kit (QIAGEN). DNA was extracted from plasma and buffy coat using Macherey-Nagel NucleoSnap and QIAGEN Blood and Tissue kits, respectively. DNA isolated from both FFPE samples and buffy coat were fragmented by sonication to 150 bp using a Covaris E220 prior to library preparation.

### Whole Exome Sequencing (WES)

Sequencing libraries were prepared using 100-500 ng of cell free DNA (cfDNA) or sonicated genomic DNA using KAPA Hyper-Prep Kit (KAPA Biosystems) with Agilent SureSelect XT Target Enrichment System and Human All Exon V5 capture baits (Agilent Technologies). Next generation sequencing was carried out using the Illumina NextSeq500 or HiSeq 2500 platform with 2×79-144 cycles by the OHSU Massively Parallel Sequencing Shared Resource to an average depth of 100X per library replicate. For Bx3 and Bx4 only, DNA isolated from both FFPE samples and buffy coat were submitted to Tempus Labs Inc. for whole exome sequencing with the Tempus xE assay.

Somatic mutation calling: sequence read FastQ data files were aligned to the UCSC hg19 human genome build using BWA MEM (0.7.12, GATK, Broad Institute) followed by marking duplicate reads (Mark Duplicates) and base recalibration (BQSR).^69,70^ Bam files for replicate libraries were merged and somatic variants were called using MuTect2 (4.0.4.0 GATK, Broad Institute) between tumor or cfDNA and the patient’s matched normal from buffy coat.^71^ A panel of normal (PON) and the gnomAD (release 2.0.1; https://gnomad.broadinstitute.org/) germline reference resource were used to filter out technical sequencing artifacts and common polymorphisms, respectively. All analysis tools were run using an OHSU Galaxy instance (v17.09).^7^

Phylogenetic and Clonal Analysis: Mutect2 and mpileup were used to call or detect presence of variants across all samples.^72^ Only sequence reads with base quality greater than 20 and mapping quality greater than 30 were used for mpileup. Variants with VAF lower than 5% or depth lower than 30 reads were filtered. The R package ape was used for phylogenetic analysis.^73^ A binary table of variants present across all tumor samples was generated as input. Genetic distance was estimated using the dist.gene function with the pairwise method. Minimum Evolution (ME) fit with ordinary least-squares (OLS) using the FastME function was used to reconstruct the phylogeny.

Copy Number Analysis: Copy number analysis was performed with CNVkit (v0.9.4a0) using the tumor/ctDNA aligned reads (BAM) and a pooled normal reference.^74^ On- and off- target read depths from each sample were median-centered log2 normalized, followed by GC bias correction and repeat masking. Tumor copy ratios were estimated by subtracting the log2-normallized depths for each bin. Corrected copy ratio profiles were segmented using circular binary segmentation (CBS). Tumor purity estimates were then used to call each segment’s absolute integer copy number.

### Dual Index Degenerate Adaptor Sequencing (DIDA-Seq)

Bait Design: Single nucleotide variants (SNVs) were filtered by frequency (>5% in the tumor/cfDNA and <2% in the matched normal) and depth (>30x in the tumor/cfDNA and >15x in the matched normal). A set of 20-40 SNVs were then hand-curated and chosen per tumor tissue sample based on clonality and potential functional impact for bait design.

Library Preparation: We chose 53 mutations representative of all four classes of origin, both clonal and subclonal, functional and non-functional, to monitor longitudinal blood draws for the presence of tumor-derived circulating tumor DNA (ctDNA). DIDA-Seq error-correction libraries were created using the Kapa Biosystems Hyper Prep kit using at least 30 ng of cell-free DNA as input as previously described, using a single over-night capture incubation instead of two incubations.^10^ Samples were sequenced on either the Illumina HiSeq 2500, paired-end 100 bp with dual 14-bp indexing cycles (highcapacity, rapid run mode) or the Illumina NextSeq 500, paired-end 75 bp with dual 14-bp indexing cycles (high-capacity, 150-cycle kit).

Error-Correction, Bait Evaluation, and Variant Analysis: The pipeline for analyzing DIDA-Seq data was based on the duplex sequencing pipeline developed in the L. Loeb laboratory (University of Washington) with substantial modification to be compatible with our data.^75^ The DIDA-Seq computational pipeline was implemented as previously described, and the variant allele frequency was determined for each mutation at each time point.^10^ Each panel of baits corresponding to variants found in a given tissue or set of tissues (primary, Bx1, and Bx2) was evaluated using unrelated patient cfDNA samples (negative controls). We sequenced each library to an average post-error correction depth of 4,000-15,000X coverage per site-of-interest and determined the variant allele frequency (VAF). We compared the mutation-specific VAF in the patient’s plasma to the VAF of the same site in the set of pooled negative controls (sequenced to an average post error-correction depth of 100,000X per site, giving an average error rate of 1 in 30,000 reads). A p-value was generated for each site and sites aggregated by panel using the overlap coefficient of the beta distributions between the VAF in the sample and VAF in the negative controls.^76^ Any site with greater than 1% VAF in the negative controls was omitted from further evaluation. Data points having a p-value greater than 0.05 were considered not statistically different from the negative controls, effectively determining our lower limit of detection given the individual or aggregated sites’ sequencing depth at each time point.

### Low-Pass Whole Genome Sequencing (LP-WGS)

LP-WGS libraries were prepared with 50 ng of sonicated tumor DNA (extracted from FFPE as described above) and patient-matched buffy coat DNA using KAPA Hyper-Prep Kit (KAPA Biosystems) with Illumina single index Tru-Seq adaptors (idtdna.com) and sequenced on the Novaseq S4 platform (Illumina) to 0.9X mean coverage. Fastq files were aligned using BWA-MEM as described above, and copy number alterations were called using the IchorCNA software package (https://github.com/broadinstitute/ichorCNA).^77^

### RNA Sequencing and Transcriptomic Analysis

Library construction and sequencing: RNA was extracted from macro-dissected, tumor-rich regions of FFPE at the CLIA-certified/CAP-accredited OHSU Knight Diagnostic Laboratories. Sequencing libraries are constructed with the TruSeq RNA Exome Library Prep Kit followed by sequencing on Illumina NextSeq500. A Universal Human Reference (UHR) (Chem-Agilent, Catalog #740000) was sequenced with each batch of samples to allow for assessment and removal of technical artifacts (due to, e.g., library preparation).

Gene Quantification: Transcript quantification followed the methods described by Tatlow and Piccolo (Tatlow and Piccolo, 2016). Briefly, the raw sequence reads were quality trimmed using Trim Galore (https://www.bioinformatics.babraham.ac.uk/projects/trim_galore/) followed by pseudo-alignment and transcription quantification using Kallisto with GENCODE reference transcriptome (version 24). Transcript level expression is aggregated to gene level abundance using the R package tximport yielding expression values for 60554 Ensembl genes.^78^

Batch Correction: Genes were filtered based on a minimum of 3 transcripts per million (TPM) in at least 3 of 48 samples, which included 29 ER+ metastatic breast cancer samples and 19 UHR samples. The filtered gene expression matrix (16,364 genes) was batch corrected by removing unwanted variation (RUV).^79,80^ RUV correction uses factor analysis to identify the factors of unwanted variation observed in the UHR batch control and corrects for them across all samples. RUV was applied by removing 1 factor (k) using the 5% of genes with the lowest standard deviation. In addition to intra-cohort batch correct, the patient samples were batch adjusted for analyses comparing to the TCGA BRCA.^81^ The TCGA BRCA gene expression matrix was filtered to samples with a Luminal (A or B) molecular subtype and joined with the RUV corrected patient sample gene expression. The combined matrix was log transformed, filtered to genes with a minimum of 3 log2 TPMs in at least 3 samples, and batched corrected using ComBat with TCGA samples set as the reference.^82^

Molecular Subtype Signature: The PAM50 subtype gene signature was used to classify samples into the intrinsic molecular subtypes.^83^ A cohort of 20 ER+ and 20 ER- samples was used as the background for classifying the patient samples’ subtypes. The gene expression matrix using these 40 samples and the patient samples was mean centered and correlated (Spearman) to the pre-defined centroids based from Parker et al. The samples were assigned to the molecular subtype with the highest Spearman correlation to the subtype centroid.

Pathway enrichment analysis: Gene set variation analysis (GSVA) was used to estimate pathway enrichment of the 1) MSigDB Cancer Hallmark Pathways (50 gene sets), 2) All MSigDB Pathways (∼20K gene sets), and 3) Reactome Pathway Database (∼2K gene sets); https://bioconductor.org/packages/release/bioc/html/GSVA.html.^21,22,84^ GSVA used a Gaussian kernel for estimating the cumulative density function and the enrichment statistic was calculated as the difference between the maximum positive/negative random walk deviations. This analysis was applied to the RUV/ComBat adjusted log2 gene expression matrix that included both TCGA BRCA Luminal Samples and the patient samples.

### Transcriptional regulator networks

Regulatory pathway and molecular interactions network: The regulatory network used to generate enrichment signatures is derived from the aggregation of publicly available molecular interactions and biological pathway databases provided by the Pathway Commons (PC) resource.^23^ The aggregated data is represented in the standard Biological Pathway Exchange (BioPAX) language and provides the most complete and rich representation of the biological network models stored in PC. These complex biochemical reactions were reduced to pairwise relationships using rules to generate a Simple Interaction Format (SIF) representation of BioPAX interactions. The reduction of BioPAX interactions to the SIF allows for the representation of pairwise molecular interactions in the context of specific binary relationships. The feature space of the SIF regulatory network was restricted to primary and secondary downstream interactions for genes within Pathway Commons. The regulatory network was then reduced to edges that are associated with the binary relationship “controls-expression-of”, defined as any reaction where the first protein controls a conversion or a template reaction that changes the expression of the second protein.

Network weight assignment: Weights are assigned to the protein-protein edges within the graph for each regulator-target pair within the regulatory network that is represented in the expression data set. These weights are derived from the integration of an F-test statistic to capture linear dependency and the Spearman rank-order correlation coefficient for a given regulator-target pair.

Regulon enrichment: This method leverages pathway information and gene expression data to produce regulon-based protein activity scores. Our method tests for positional shifts in experimental-evidence supported networks consisting of transcription factors and their downstream signaling pathways when projected onto a rank-sorted gene-expression signature. The gene-expression signature is derived by comparing all features to the median expression level of all samples considered within the data-set. After weights have been assigned to the regulatory network, the positive and negative edges of each regulator are rank ordered. The first component of the enrichment signature, the local delta concordance signature, is derived by capturing the concordance between the magnitude of the weight assigned to a particular edge and the ranked position of that edge. The features associated with activation, positive edges within the regulatory network, are monotonically ranked from most lowly to highly expressed in the restricted feature space, where the features that are repressed are ranked by a monotonically decreasing function. This component of the signature considers positive and negative edges independently, which captures support for an enrichment signature even if one of the edge groups is underrepresented in the network graph. The second component of the enrichment signature, the local enrichment signature, captures positional shifts in the local gene ranked signature and integrates those shifts with the weights assigned to overlapping features for a given regulon and the expression data set. The last component of the enrichment signature considers the entire feature space and projects the rank-sorted local signature onto this global ranked feature space. We derive our global enrichment signature from this projection for each regulator we consider. We use the median of robust quantile-transformed ranked positions as the enrichment scores for both the local enrichment and global enrichment signatures. We then integrate these three individual signatures together, which allows us to capture differences between individual regulator signatures within the context of an individual patient as well as at a cohort level.

### Reverse Phase Protein Arrays

Protein extracts from tumor samples were analyzed as previously described.^85,86^ In order to scale the protein expression values, the RPPA data from the patient samples was merged within the TCGA breast cancer RPPA dataset, using the replicate-based normalization method.^87^ The protein expression values were then z-scored by using the median and standard deviation, and a heat-map was generated from the treated and untreated samples, using Rank-Sum ordering of the proteins fold change. The heat map was produced using publicly available Cluster 3.0 and TreeView software.

Pathway Analysis: All pathway predictors have been previously described.^85^ Proteins used as predictors of the different pathways are listed in Table S2. To determine a pathway score, for each sample all positively associated predictors were summed minus the predictors that are negatively associated with the pathway. The total was then divided by the numbers of predictors in the pathway. To generate the pathway scores histograms, the distribution of each TCGA samples subtype was plotted and the value of the patient pre- and post-treatment sample was added to the histograms.

### Intracellular Signaling Protein Panel

The Nanostring 3D Vantage Solid Tumor Panel is comprised of 27 antibodies, including 13 targeting phosphorylated proteins, specifically designed to interrogate the MAPK and PI3K/mTOR signaling pathways.^13^ This multiplexed panel allows for the simultaneous quantification of multiple proteins from a single section of FFPE tissue. Four micrometer sections of FFPE cancer cell lines (controls) or tumor biopsy tissue were subjected to citrate-based antigen retrieval and incubated overnight with the cocktail of oligo-tagged antibodies. After washing, the oligo-tags were released by UV light (3 minutes on a UV lightbox) and quantified using the Nanostring nCounter system. A set of 6 FFPE cancer cell lines were selected as positive controls and included on every run to assess antibody and control performance and to correct for batch effects. Batch correction was preformed using Removal of Unwanted Variation (RUV) using the replicate positive controls to estimate the factors associated with batch effect.^88^ RUV parameters were optimized by measuring the consistency of replicate controls and careful evaluation of outliers to ensure validity of results. This assay was validated for clinical use in the Knight Diagnostic Laboratories at OHSU as the GeneTrails^®^ Intracellular Signaling Protein Panel.

### Integrative Analyses

Multi-omic integrated pathway analysis: CausalPath (https://github.com/PathwayAndDataAnalysis/causalpath; commit 9f8d6f8) was used for integrated pathway analysis of protein, phosphoprotein, gene abundance, and transcriptional regulator activity.^24^ Briefly, CausalPath is a hypothesis generating tool that uses literature-grounded interactions from Pathway Commons to produce a graphical representation of causal relationships that are consistent with patterns in a multi-omic datasets.^24^ This integrative approach allows for holistic evaluation of signaling networks and pathway activity across longitudinal biopsies. The CausalPath analysis used the log fold change of total and phosphoprotein (RPPA) and gene expression from Bx1 and Bx2 with the following parameters: threshold-for-data-significance = 0.3 for RNA, protein, and phosphoprotein, value-transformation = max, calculate-network-significance = true, permutations-for-significance = 10,000, color-saturation-value = 2.5, data-type-for-expressional-targets = rna and protein, show-all-genes-with-proteomic-data = true. The resulting network was pruned to include the neighborhoods encompassing MTOR, AKT, MUC1, STAT3, MYC, and E2F1 to highlight biologically interesting patterns discussed in the text. For additional depth, the difference in transcriptional regulon enrichment activity between Bx2 and Bx1 was mapped to and overlaid on the pruned CausalPath network.

Integrated Heatmap: The gene, protein, phosphoprotein abundances, and transcriptional regulon enrichment activities were integrated into a single heatmap. Each data type was independently scaled to −1 to 1 with the exception of protein/phosphoprotein, which were scaled together. Fold change of biopsy 2 to biopsy 1 was calculated for each scaled feature and represented as a heatmap grouped by pathway categories of interest.

### Multiplex Immunohistochemistry

Immunohistochemical staining: Glass mounted FFPE tissue sections (5 μm) were baked at 60°C for 60 minutes, deparaffinized with xylene, and rehydrated in serially graded alcohols, then place in distilled water. Slides were stained with hematoxylin (Dako, S3301) for 1 minute, mounted with 1x TBST buffer (Boston Bioproducts, IBB-181R), coverslipped with Signature Series Cover Glass (Thermo Scientific, 12460S), and subjected to whole slide digital scanning at 20x magnification using an Aperio ImageScope (Leica Biosystems). Slides were de-coverslipped with 1 min of agitation in TBST and subjected to heat-mediated antigen retrieval in 1x pH 6.0 citrate buffer (Biogenex Laboratories, HK0809K) for 20 min at 95°C followed by blocking of endogenous peroxidase activity (Dako, S2003, per manufacturer’s instructions). Slides were then subjected to 12 cycles of multiplex immunohistochemistry (mIHC), each cycle consisted of either 1 or 2 rounds of IHC. Each round consisted of immunodetection (primary antibody, HRP-linked secondary antibody, HRP-mediated development of AEC chromogen), and whole slide scanning. Citrate antigen retrieval was used between cycles to remove primary antibodies, and HRP inactivation was used between rounds (Dako, S2003, per manufacturer’s instructions) to eliminate HRP carry-over as described previously.^14,15^ Several antibody panels (and variations of) were utilized for the current study (Table S2). Where IHC and chromogenic staining did not pass QC, they were not included in analysis: e.g., PD-L1 and CSF1R on the myeloid panel, and CD68 and ICOS on the functional panel(s). Several antibodies were not common across all or some panels, thus not included in results: IDO on functional panel (Bx1), Tryptase on myeloid panel (Primary, Bx1 and Bx2), RORyT and GATA3 on the lymphoid panel (primary, Bx1, and Bx2), and CCR2, HLA class-I, CD169, CD11b, and CD11c on the discovery panel (23 antibodies) (Bx3).

Image analysis pipeline: Regions of interest (ROIs) were selected from hematoxylin-stained images reflecting histopathologic regions containing either primary or metastatic tumor foci via analysis in ImageScope (Leica). Digitally-acquired images were registered in MatLab (MathWorks) utilizing the SURF algorithm in the Computer Vision Toolbox. Nuclear segmentation and color deconvolution were performed using an in-house FIJI macro (ImageJ, NIH) for segmenting hematoxylin only stained tissue. In short, preprocessing to isolate signal and remove background was performed, then nuclear objects were identified by watershed and standard image processing (erosion, dilation, and noise removal). AEC chromogenic signal was extracted by converting images from RGB to CMYK in ImageJ with the NIH plugin RGB_to_CMYK. The contrast of AEC chromogen intensities on a 0-255 scale in the yellow channel, as compared to RGB or the built-in AEC deconvolution vector, utilized the full intensity scale without a threshold. First, each channel was normalized by dividing all pixels in each image by the max intensity of that image to rescale image intensity values to a range 0 - 1. Next, rescaled isolated signal from each stain was quantified for every indexed nuclear object by a pipeline created in Cell Profiler 3.1.5 (Broad Institute). Image cytometry hierarchical gating was performed in FCS Express Image Cytometry RUO 6.1.4 (DeNovo Software) to quantify distinct populations of cells with spatial context.

### Cyclic Immunofluorescence

Immunofluorescence analyses of tumor tissue: FFPE human tissues were sectioned at 4 μm and mounted on adhesive slides (Mercedes Medical, TNR WHT45AD). The slides were baked overnight in an oven at 55 °C (Robbin Scientific, Model 1000) and an additional 30 minutes at 65 °C (Clinical Scientific Equipment, NO. 100). Tissues were deparaffinized with xylene and rehydrated with graded ethanol baths. Two step antigen retrieval was performed in the Decloaking Chamber (Biocare Medical) using the following settings: set point 1 (SP1), 125 °C, 30 seconds; SP2: 90 °C, 30 seconds; SP limit: 10 °C. Slides were further incubated in hot pH 9 buffer for 15 minutes. Slides were then washed in two brief changes of diH_2_O (∼2 seconds) and once for 5 minutes in 1x phosphate buffered saline (PBS), pH 7.4 (Fisher, BP39920). Sections were blocked in 10% normal goat serum (NGS, Vector S-1000), 1% bovine serum albumin (BSA, Sigma A7906) in PBS for 30 minutes at 20 °C in a humid chamber, followed by PBS washes. Primary antibodies (Table S2) were diluted in 5% NGS, 1% BSA in 1x PBS and applied overnight at 4° C in a humid chamber, covered with plastic coverslips (IHC World, IW-2601). Following overnight incubation, tissues were washed 3 x 10 min in 1x PBS. Coverslips (Corning; 2980-243 or 2980-245) were mounted in Slowfade Gold plus DAPI mounting media (Life Technologies, S36938).

Fluorescence Microscopy: Fluorescently stained slides were scanned on the Zeiss AxioScan.Z1 (Zeiss, Germany) with a Colibri 7 light source (Zeiss). The filter cubes used for image collection were DAPI (Zeiss 96 HE), Alexa Fluor 488 (AF488, Zeiss 38 HE), AF555 (Zeiss 43 HE), AF647 (Zeiss 50), and AF750 (Chroma 49007 ET Cy7). The exposure time was determined individually for each slide and stain to ensure good dynamic range but not saturation. Full tissue scans were taken with the 20x objective (Plan-Apochromat 0.8NA WD=0.55, Zeiss), and stitching was performed in Zen Blue image acquisition software (Zeiss).

Quenching Fluorescence Signal: After successful scanning, slides were soaked in 1x PBS for 10 – 30 minutes in a glass Coplin jar, waiting until the glass coverslip slid off without agitation. Quenching solution containing 20 mM sodium hydroxide (NaOH) and 3% hydrogen peroxide (H_2_O_2_) in 1x PBS was freshly prepared from stock solutions of 5 M NaOH and 30% H_2_O_2_, and each slide placed in 10 ml quenching solution. Slides were quenched under incandescent light, for 30 minutes for FFPE tissue slides. Slides were then removed from the chamber with forceps and washed 3 x 2 min in 1x PBS. The next round of primary antibodies was applied, diluted in blocking buffer as previously described, and imaging and quenching were repeated over ten rounds for FFPE tissue slides.

Cyclic IF quantification and analysis: Each image acquired during the Cyclic IF assay was registered based on DAPI features acquired from each round of staining.^89^ Cellpose, a generalist algorithm for cellular segmentation, was used to generate nuclear and cell segmentation masks with a pre-trained neural network classifier.^90^ Extracted single-cell features included centroids and mean intensity of each marker from its biologically-relevant segmentation mask (e.g., Ecad_Cytoplasm, Ki67_Nuclei). The last round DAPI image was used to filter out cells lost during each round of Cyclic IF staining.

For cell type determination and composition analysis, single cell mean intensities from each biopsy were batch corrected using the ComBat algorithm.^91^ ComBat was used to adjust the mean and variance of fluorescence intensity on control tissue-microarrays (TMAs) that were stained with each biopsy, and the same adjustments were applied to the corresponding biopsies. Eighteen markers were selected for clustering; some markers were excluded due to tissue loss in the TMA controls. Principal component analysis was performed with scanpy to reduce dimensionality, and Umap was run on the top 17 principal components to calculate a nearest neighbor graph based on the 30 nearest neighbors.^92,93^ Leiden clustering was performed on the nearest neighbor graph to define clustering-based cell types.^94^ The Leiden clustering resolution of 0.5 was selected based on appropriate clustering of technical replicates in the control TMAs.

Immune, endothelial, and stromal cells were identified by manual thresholding and gating. Endothelial cells were defined as CD31+, immune cells were either CD45+ or CD68+ and CD31-, and stromal cells were cytokeratin-, E-cadherin-, CD31-, CD45-, and CD68-. Tumor was defined as cytokeratin+, and proliferating cells were Ki67+. Cell segmentation borders of manually defined cell types were visualized on the images using napari (https://zenodo.org/record/4046812/export/hx).

To calculate distance to extracellular matrix proteins, a threshold was applied to create a pixel mask of positive staining. The distance from each nuclear centroid to the nearest mask pixel was measured. Cells were grouped into bins of 0-25 microns, 25-50 microns, and 50-75 microns from the mask, and the intensity distributions were compared using ANOVA.

### Scanning Electron Microscopy

Tissue for scanning electron microscopy (SEM) was collected at the time of biopsy and placed into SEM-specific fixative (2.5% paraformaldehyde, 2.5% glutaraldehyde in 0.1M sodium cacodylate buffer) as soon as possible for both pre-treatment and on-treatment biopsies. No tissue was collected for SEM from the third biopsy due to it originating from bone and decalcification protocols cannot yet reliably preserve ultrastructure for electron microscopy. Tissues were stored at 4°C in fixative and can remain that way nearly indefinitely.

Tissue samples were prepared for SEM by implementing a post-fixation heavy metal infiltration followed by epoxy-resin embedding (Epon 812). Heavy metal staining using osmium tetroxide, uranyl acetate, and lead aspartate provided contrast for imaging by dissociating the metals and allowing them to bind to lipids and proteins within cellular membranes and organelles. After staining and resin embedding, polymerized blocks were mounted directly to SEM pin-style stubs and trimmed to create a flat surface using a Leica UC7 ultramicrotome equipped with Diatome diamond knives. Mounted blocks were conductively coated with 8-nm carbon using a Leica ACE600 coater.

Two-dimensional large-format SEM maps were collected on trimmed block faces using a FEI Helios NanoLab G3 DualBeam™ focused ion beam-scanning electron microscope (FIB-SEM) equipped with the Thermo Scientific Maps™ software package. Using this software for automation, hundreds of tiled images were collected over the entire block surface and stitched together, creating a pyramidal viewing architecture that provides observations starting at the millimeter-scale and zooms all the way down to 4-nm/pixel spatial resolution. Imaging conditions were 3 keV, 200-400 pA, 4-mm working distance, and 3 μs dwell time using the concentric backscatter detector (CBS). A script developed in-house converts these large maps into ome.tiff format for web-based viewing.

Regions of interest for three-dimensional electron microscopy (3DEM) were selected from the high-resolution maps. Three separate 3DEM datasets collected using FIB-SEM technology were generated using vendor-specific automated serial-sectioning software: two high-resolution, small volumes (4-nm/voxel, 25 x 20 x 6-10 μm^3^) on each respective biopsy and one lower resolution, larger volume (10-nm/voxel, 48 x 48 x 17 μm^3^). The high-resolution image stacks were collected using the aforementioned Helios FIB-SEM with the same electron beam conditions and the In-Column Detector (ICD). The large volume was collected from the pre-treatment biopsy using a Zeiss Crossbeam 550 FIB-SEM using 1.5 keV, 1.0 nA, 5-mm working distance, 1.6 μs dwell time, and the Energy-Selective Backscatter (EsB) detector. Segmentation of image stacks was performed manually in-house and by a CloudFactory managed workforce. Deep learning models developed in-house were utilized for nucleus and nucleoli segmentation on the high-resolution image stacks.

A more detailed description of the fixation procedure, sample preparation protocol, and imaging workflow can be found in a recently published open access book chapter^19^ and via protocols.io:

**“**Sample fixation of biopsy tissue for Electron Microscopy (EM)**”** (dx.doi.org/10.17504/protocols.io.4bigske)

“Post-Fixation Heavy Metal Staining and Resin Embedding for Electron Microscopy (EM)” (dx.doi.org/10.17504/protocols.io.36vgre6)

“2D and 3D Electron Microscopy (EM) Imaging of Tissue Biopsies and Resections” (dx.doi.org/10.17504/protocols.io.bg58jy9w)

All raw data, metadata, image stacks, and segmentation files are publicly available via the HTAN data portal and viewable via OMERO using the PathViewer plug-in.

### Stochastic Optical Reconstruction Microscopy

SKBR3 cells (ATCC HTB0-30) were cultured in McCoy’s 5A medium (Thermo Fisher Scientific 16600082) supplemented with 10% FBS (Thermo Fisher Scientific 10082147). For Stochastic Optical Reconstruction Microscopy (STORM) experiments, the cells were plated in LabTek chambered coverglass (Thermo Fisher Scientific 155409) for 36 to 48 hours before labeling and imaging. To prepare for imaging, the cells were first serum starved overnight (∼16 hr); on the day of imaging, the cells were treated with 100 nM Alexa Fluor 647 conjugated trastuzumab for ∼15 min, washed with pre-warmed blank medium, and placed on the microscope stage for imaging. Next, fresh STORM imaging buffer was added at 1:1,000 v/v dilution to the medium; the buffer is PBS supplemented with 0.5 mg/mL glucose oxidase (Sigma-Aldrich, G2133–50 kU), 40 μg/mL catalase (Sigma-Aldrich, C100-50MG), and 10% D-Glucose (w/v, Fisher Chemicals D16–500); this was followed by addition of 10 mM (final concentration) mercaptoethylamine (MEA; Sigma-Aldrich, 30070). The sample was then explored at low 647 nm laser power (∼100 W/cm^2^; this avoids unnecessary loss of AF647 due to photobleaching) to identify regions of interest. EGF (Cell Signaling 8916) was diluted from a 1 mg/mL stock in PBS to 10 mg/mL and then added to the cell culture at 1:100 v/v dilution to yield a final concentration of 10 ng/mL. Image acquisition was initiated right after adding EGF, as described below. Throughout the imaging process, the cells were kept in an on-stage incubator (TokaiHit) at 37°C with 5% CO_2_.

The STORM microscope setup was the same as described previously.^95^ Briefly, a custom single-molecule fluorescence imaging setup was built on a Nikon Ti-U microscope frame, with other essential components including an objective lens with high numerical aperture (Nikon 60x oil, TIRF, NA = 1.49), a 647 nm laser (Coherent OBIS, max output = 140 mW; for exciting and converting AF647 into a dark state), a 405 nm laser (Coherent CUBE; for converting AF647 to fluorescent on-state), and an EM-CCD (Evolve 512 Delta, Photometrics), as well as other components including dichroic mirrors and emission filters. Image acquisition was performed using micromanager with an EM-CCD gain setting typically set at 300 and the frame acquisition time 8 ms (possible by selecting a small region of interest).^96^ Typical power densities for the 647 nm and the 405 nm lasers were 1-2 kW/cm^2^ and 1-20 W/cm^2^, respectively. Raw STORM images were processed and reconstructed using custom Matlab scripts.^97^

Alexa Fluor 647 conjugated trastuzumab was prepared using Alexa Fluor 647 NHS-ester (Thermo Fisher Scientific A37566) and purified according to manufacturer recommended procedures; the final dye to antibody conjugation ratio was measured to be around 2:1 using a UV-Vis spectrometer.

## Supplemental Information

**Figure S1.**
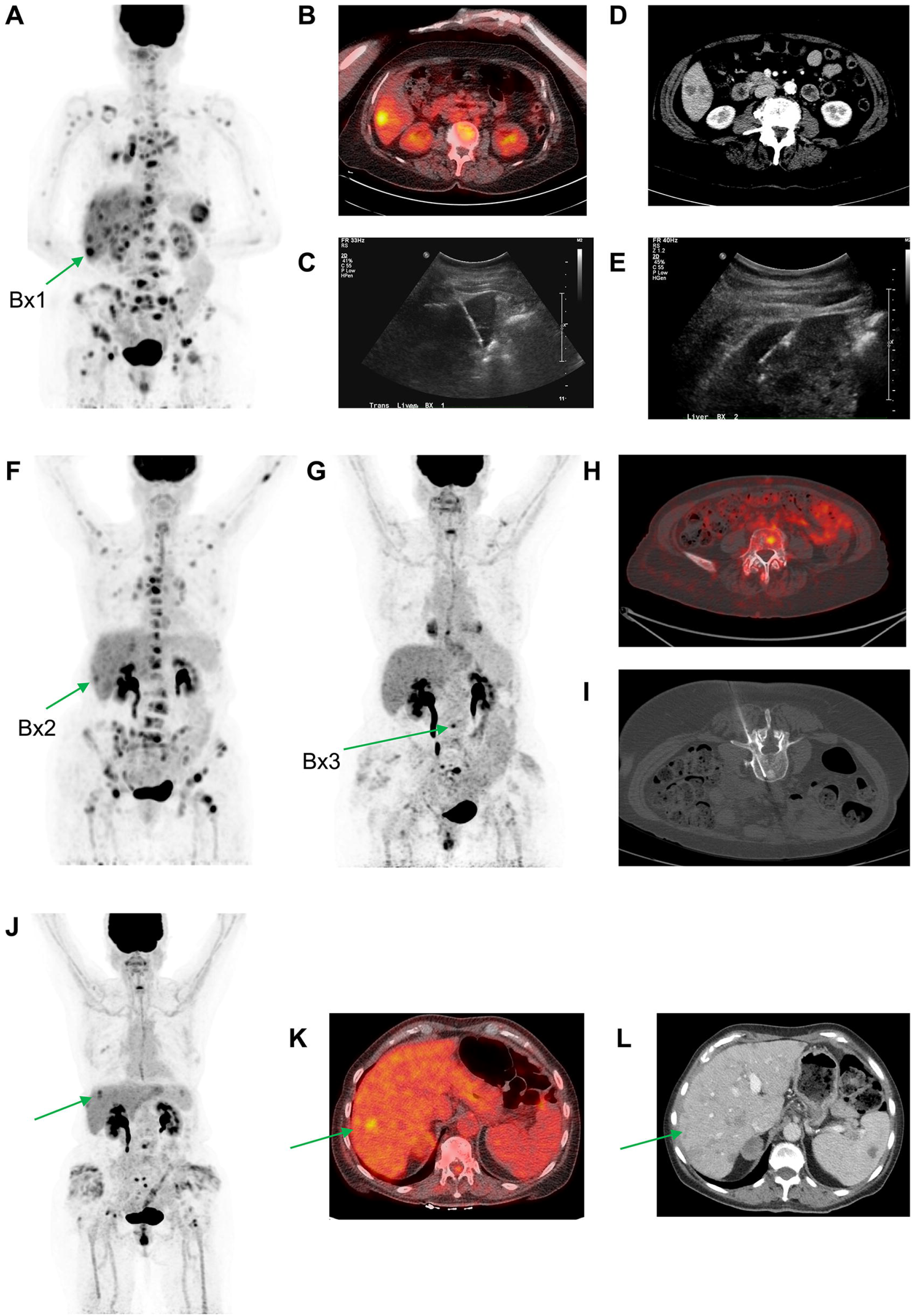
Related to Figure 2 (A) Maximum intensity projection (MIP) FDG-PET from the beginning of Phase 1 treatment demonstrates multifocal FDG avid disease throughout the mediastinum, liver, spleen, and skeleton, including target of Bx1 (arrow). (B) Axial FDG-PET (from same timepoint as in A) superimposed on CT showing FDG avid segment 6 liver lesion targeted by Bx1. (C) Ultrasound image from Bx1. (D) Axial contrast enhanced CT taken just before Bx2 demonstrates interval growth of a separate segment 5,6 liver lesion subsequently targeted by Bx2. (E) Ultrasound image from Bx2. (F) FDG-PET MIP image from the beginning of Phase 3 demonstrates a decrease in FDG avid lesions. Bx2 targeted lesion indicated (arrow). (G) FDG-PET taken in Phase 3 one month before Bx3 demonstrates new FDG avid lesions in the L4 vertebral body, including the Bx3 targeted lesion (arrow). (H) Axial FDG-PET (from same timepoint as in G) superimposed on attenuation correction CT showing FDG avid segment targeted by Bx3. (I) CT image from Bx3 demonstrates successful biopsy of the FDG avid, lytic lesion within the L4 vertebral body. Note the patient is prone during the biopsy. (J) FDG-PET from the end of Phase 3 demonstrates continued response in most organs but a possible new progressing liver lesion subsequently targeted by Bx4 (arrow). (K) Axial FDG-PET (from same timepoint as J) superimposed on CT showing FDG avid liver lesion targeted by Bx4 (arrow). (L) Axial contrast enhanced CT taken during month 37 demonstrates nodular, heterogeneous morphology of the liver showing signs of pseudocirrhosis (arrow).

**Figure S2.**
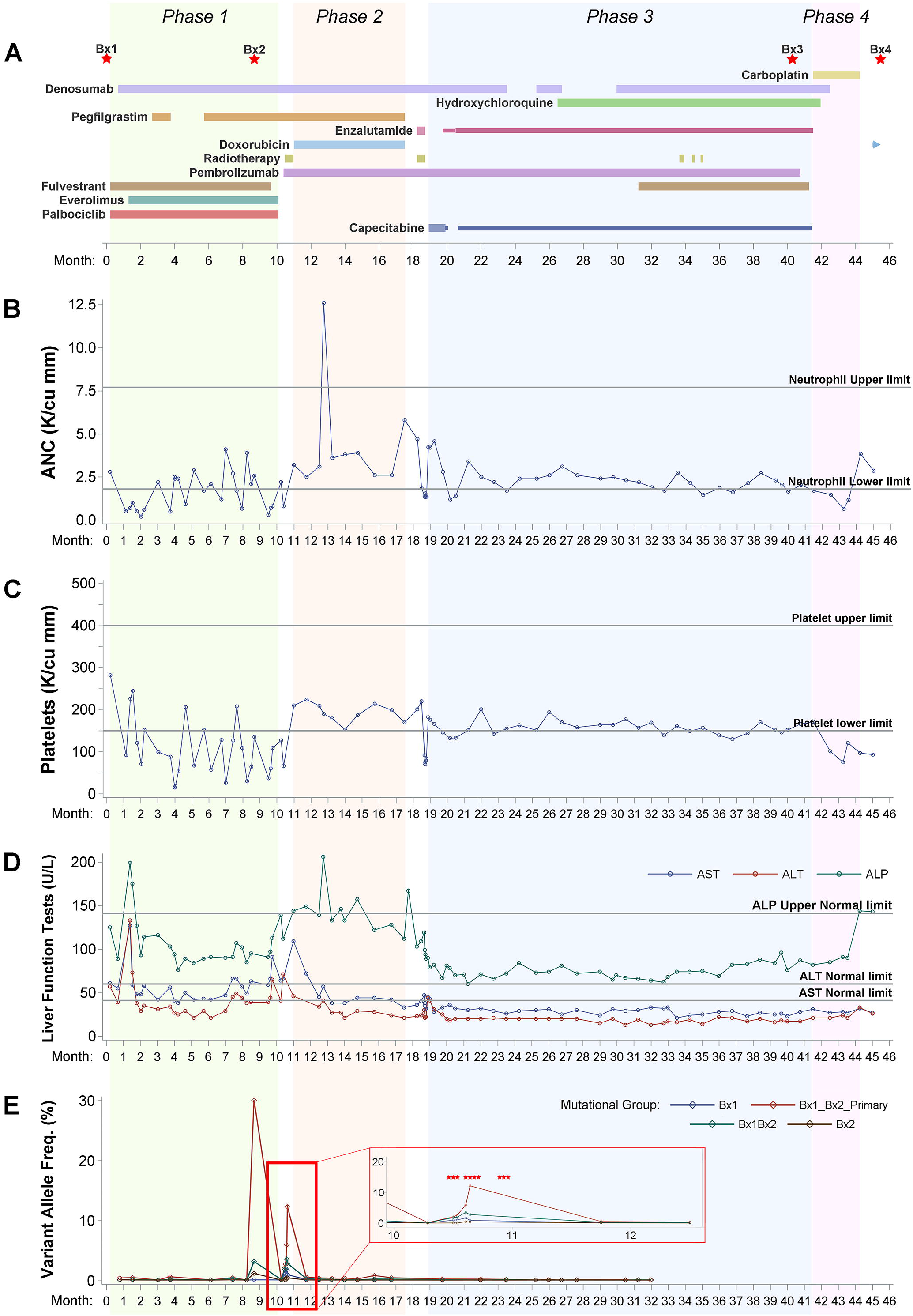
Related to Figure 2 (A) Treatment schedule and biopsy timing (red stars) over the course of four phases of treatment (green, orange, blue, and pink areas). Timeline sectioned into 28-day months. The duration and relative dose for each drug is indicated by the extent and width of a horizontal bar, respectively. Continuation of a drug after the end of Phase 4 is indicated by a right pointing arrow. (B) Clinically reported Absolute Neutrophil Count (ANC) in thousands per cubic millimeter (K/cu mm). (C) Clinically reported Platelet Count in thousands per cubic millimeter (K/cu mm). (D) Clinically reported results of liver function tests, including alkaline phosphatase (ALP), alanine aminotransferase (ALT), and aspartate aminotransferase (AST). (E) Longitudinal tracking of the average circulating tumor DNA (ctDNA) Variant Allele Frequency (VAF) of four different groups of mutations: Variants private to Bx1 (Bx1), private to Bx2 (Bx2), shared between the primary, Bx1, and Bx2 (Bx1_Bx2_Primary), and shared by Bx1 and Bx2 but not the primary (Bx1_Bx2). Red boxed inset shows expanded ctDNA VAF timeline during a course of palliative radiotherapy during month 10. Red asterisks indicate dates of individual radiation fractions given.

**Figure S3.**
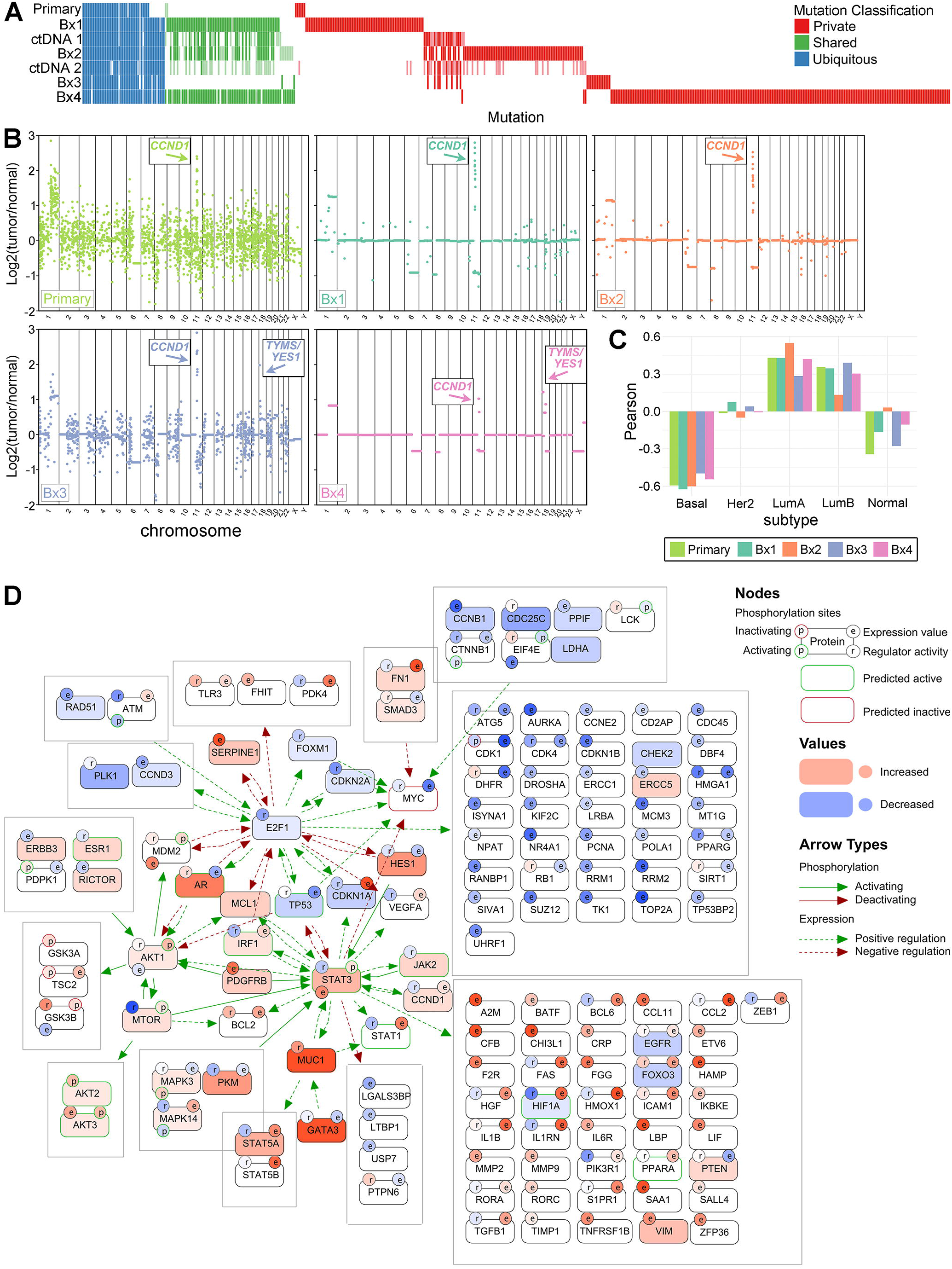
Related to Figure 3 (A) Non-silent SNVs and Indels identified from WES of tissue samples and classified as Ubiquitous (blue), Shared (green), or Private (red) (variants private to ctDNA timepoints not shown). Mutational status in each biopsy sample or in circulating tumor DNA from peripheral blood is indicated as independently called (colored), detected in at least 2 sequencing reads but not independently called (reduced opacity), or absent (white). (B) Scatter plots of genome-wide, log2 copy number profiles from WES (Primary, Bx1, Bx2, and Bx3) and LP- WGS (Bx4): Primary (yellow-green), Bx1 (green), Bx2 (orange), Bx3 (blue), and Bx4 (pink). (C) Molecular Subtype. Bar plots show the Pearson correlation of the Primary (yellow-green), Bx1 (green), Bx2 (orange), Bx3 (blue), and Bx4 (pink) samples to the PAM50 subtype centroids. (D) Integrated multi-omic pathway analysis. Pathway diagrams generated with CausalPath represent the integration of protein abundance (rectangles), phosphoprotein abundance (circle labeled ‘p’; green outline indicates activating; red outline indicates inactivating), gene expression (circle labeled ‘e’), and transcriptional regulator activity (circle labeled ‘r’), and show the change in Bx2 relative to Bx1. Networks were generated using protein/phosphoprotein abundance and gene expression, while transcriptional regulator activity was mapped on following network pruning. The red and blue fill represent higher and lower expression/activity, respectively.

**Figure S4.**
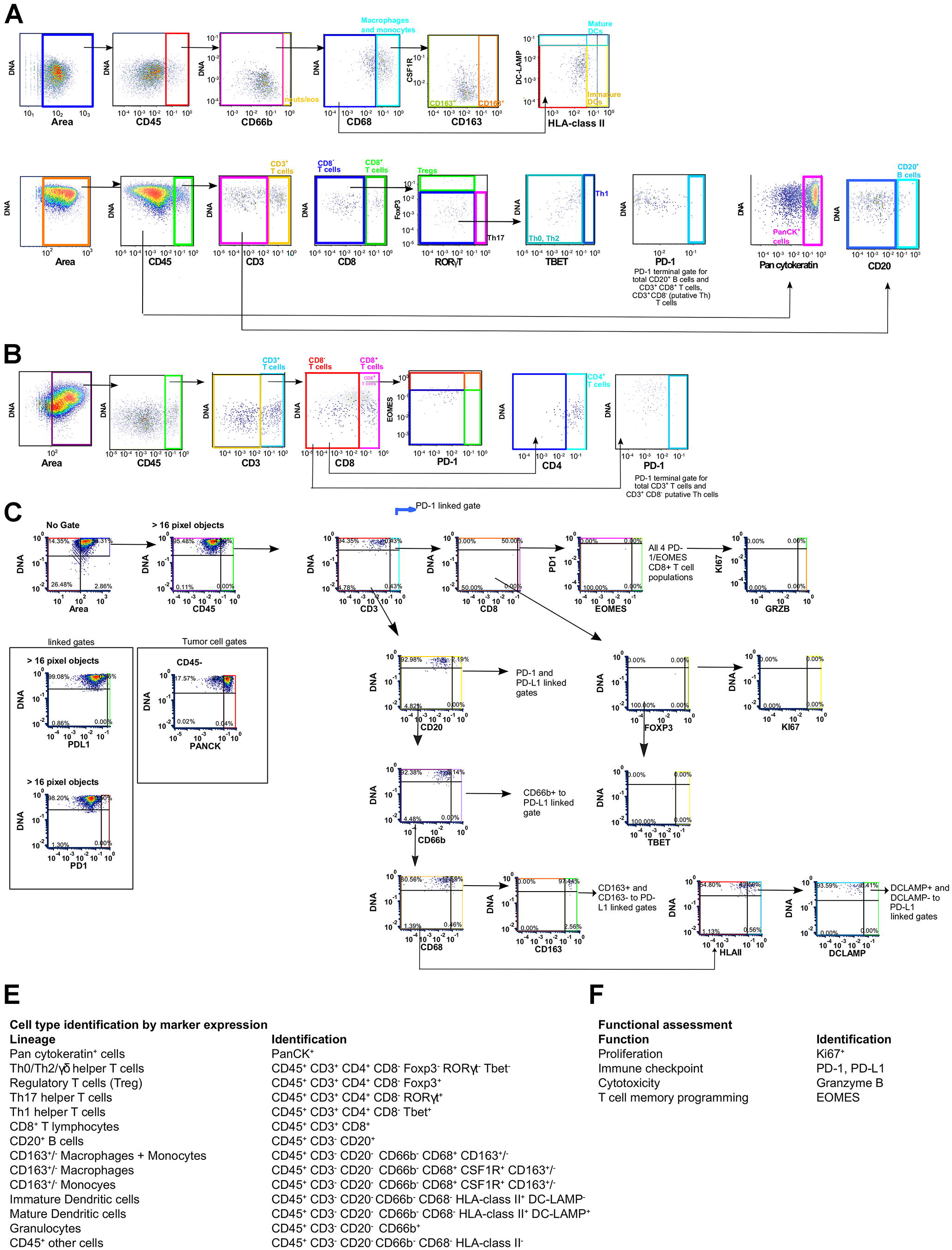
Related to Figure 4 Hierarchical gating, cell identification, and supplementary data for mIHC studies. Hierarchical gating via image cytometry for identification of cell types and functional states elaborated by mIHC for: (A) myeloid, (B) lymphoid, (C) functional, and (D) discovery panel of antibodies. (E) Classification of immune cell types and (F) functional status by marker expression assessed in hierarchical gating.

**Figure S5.**
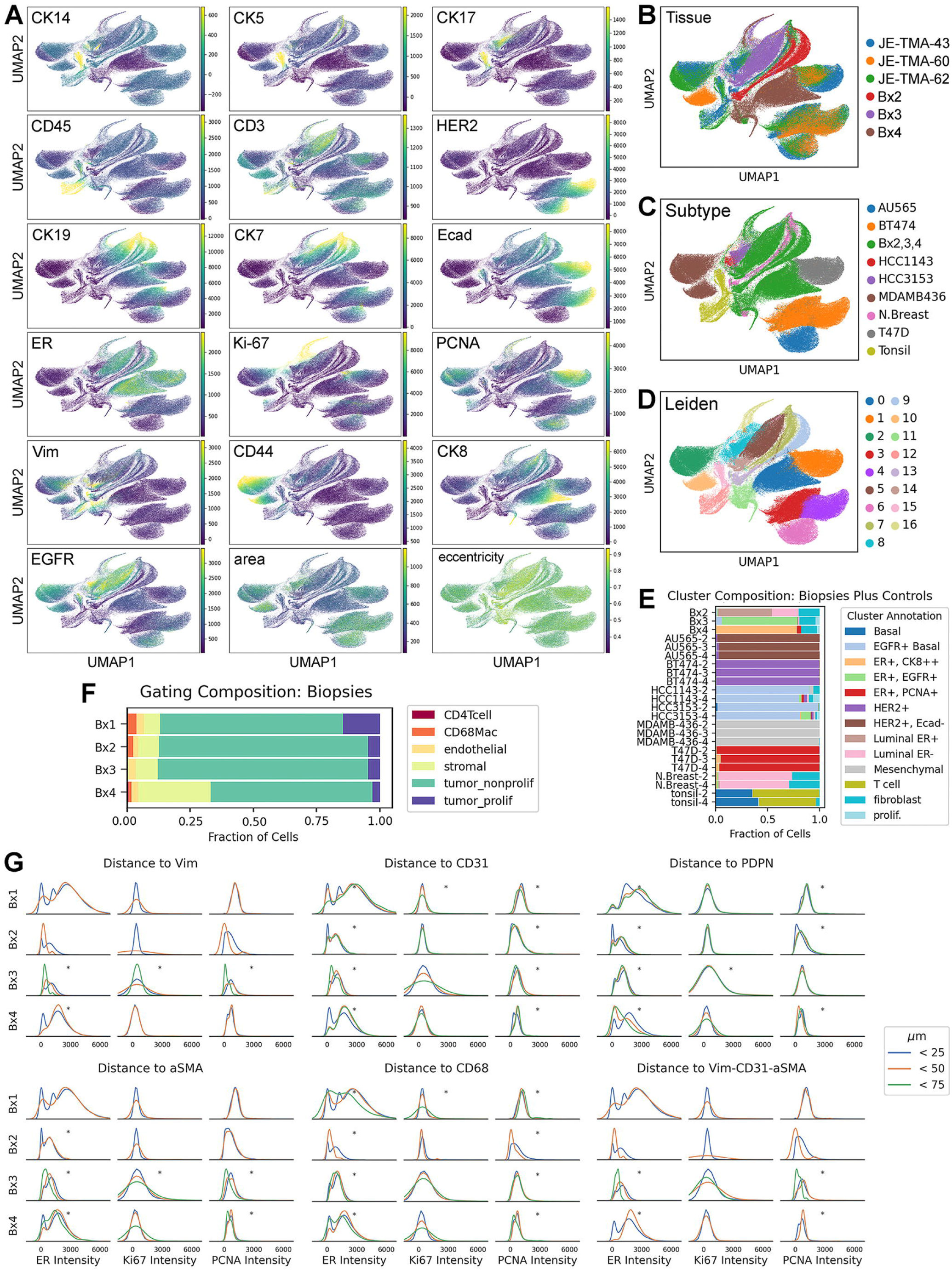
Related to Figure 5 (A) Umap projection of 18 features used for clustering, colored by feature. (B) Tissues included in cluster analysis. Controls for each biopsy are as follows, Bx2: JE-TMA-43, Bx3: JE-TMA-60, Bx4: JE-TMA-62. TMAs were used for normalization, so mixing of cells from all TMAs indicates successful normalization. (C) Umap colored by cell lines and normal tissues included in clustering. (D) Umap projection colored by Leiden clustering result. (E) Composition of biopsies and controls by annotated Leiden cluster. Number following cell line and tissue names indicates the biopsy that was paired with those controls. (F) Composition of biopsies based on manual thresholding and gating (see methods). (G) Intensity of ER, Ki67, and PCNA at 0-25, 25-50, and 50-75 μm away from various markers. Asterisks indicate significant (p < 0.001) difference in mean intensity between distances (ANOVA).

**Figure S6.**
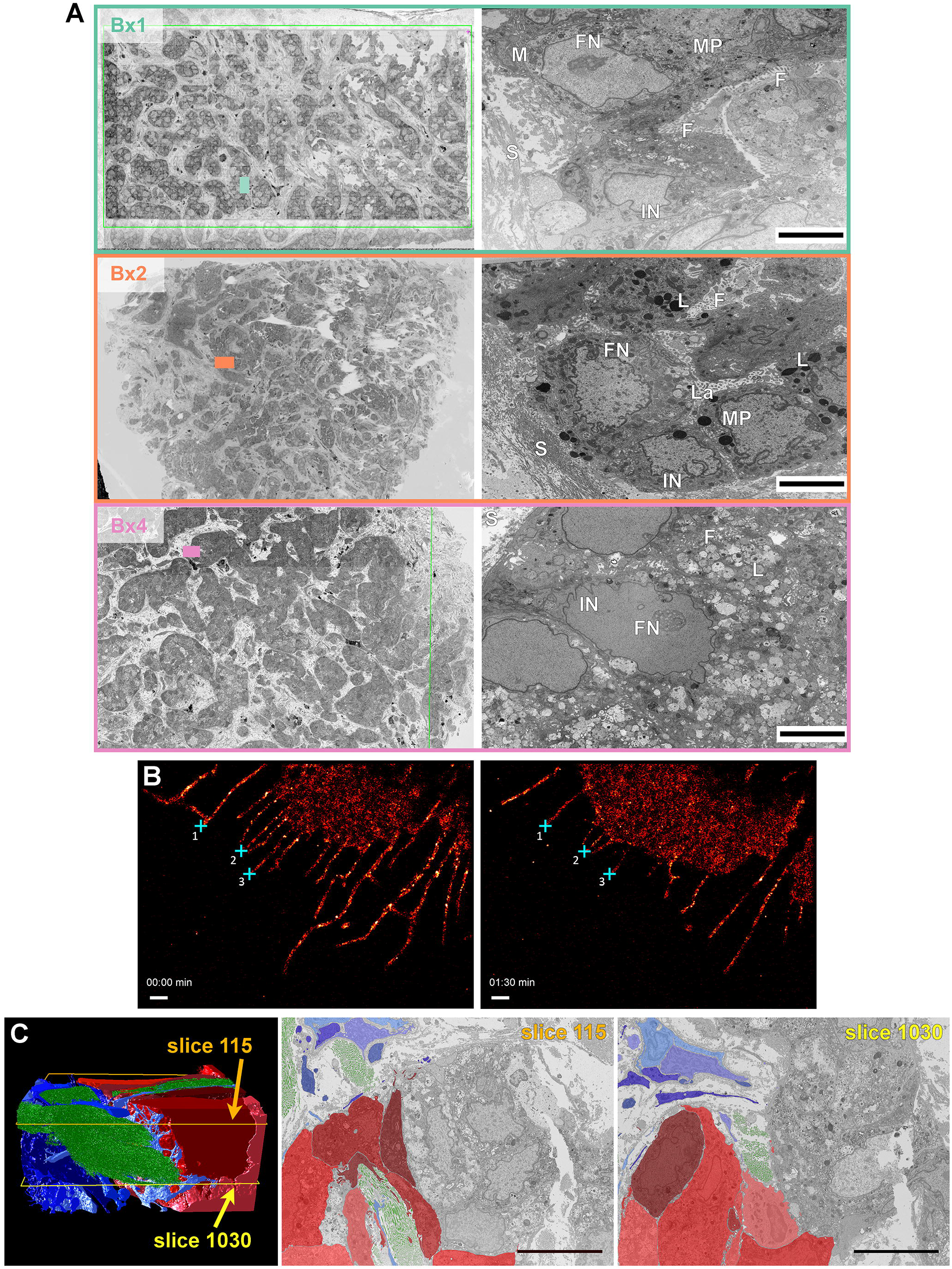
Related to Figure 6 (A) The left column shows top-down, high-resolution blockface maps collected via 2D SEM from Bx1, Bx2, and Bx4. The boxes marked on the maps measure 25 μm in the long direction and indicate where 3D FIB-SEM was collected. The right column shows the respective first slice from the FIB-SEM volumes. Ultrastructural features of interest are marked as the following: (IN) invaginated nuclei, (FN) fenestrated nucleoli, (M) mitochondria, (L) lysosome, (S) stroma, (F) filopodia, (La) lamellipodia, (MP) macropinosomes. Scale bars, 4 μm. Bx1 shows well-defined nests of tumor cells separated by thick bands of collagen. Bx2 also shows tumor cell nests, but the tissue is denser and collagen band thickness is reduced. Bx4 shows a return to thick stromal bands and clear tumor cell nest formation, but the high-resolution view on the right shows ultrastructure different to Bx1, particularly with respect to lysosomes and macropinosomes. (B) Filopodia-like protrusions (FLPs) direct EGF-induced cell movement of SKBR3 cells. Time-lapse stochastic optical reconstruction microscopy (STORM) images of SKBR3 cells labeled with Alexa Fluor 647-conjugated Herceptin, showing abundant FLPs decorated by HER2. Left image shows initial positions of the FLPs upon addition of EGF (10 ng/mL at around -30 s). Right image shows the same field of view after 1:30 min (90 s). The cyan crosses (1, 2, and 3) mark the original locations of the tips of corresponding FLPs. Scale bars, 1 mm. (C) Segmentation of any imaging modality is limited by the 2D plane being viewed. As shown by the FIB-SEM volume, the organization, number, and area of tumor cells (red), stromal cells (blue), and collagen (green), are different depending on the depth (image slice number) of the 2D plane within the tissue sample. Slices 115 and 1030 are separated by ∼9 μm. Scale bars, 10 μm.

Table S1. Related to Figure 2

Clinical metadata. Includes dates of all individual drug doses, results of serum tumor protein biomarker assays, neutrophil and platelet counts, liver function tests, CT and FDG-PET lesion measurements, and clinically reported immunohistochemical (IHC) assay results.

Table S2. Related to Figures 3-5

Exploratory analytic information. Includes gene set variation analysis scores of MSigDB databases (RNAseq); transcriptional regulator activity scores (RNAseq); pathway signature analysis scores (RPPA); pathway signature analysis weights (RPPA); antibody information and staining order for panels focused on lymphoid, myeloid, functional, and combined (discovery, mIHC); and antibodies used (CycIF).

Movie S1. Related to Figure 6

The 3D FIB-SEM volume collected at 4 nm/voxel resolution from Bx1 showing ultrastructural features at the nanoscale. Individual cell contours are rendered and illustrate the cell-cell and cell-stromal interactions.

Aberrant nuclear morphology, clustered macropinosomes, organized mitochondria, and the presence of lysosomes are all observed in the 25 x 20 x 6 μm^3^ volume.

Movie S2. Related to Figure 6

A larger 3D FIB-SEM volume (60 x 40 x 18 μm^3^) collected at 10 nm/voxel resolution of Bx1 shows tumor nest interaction with the fibroblasts in the stroma. The fibroblasts and stromal cells in blue are interacting with the red cancer cells and wrapping themselves around the nest to form a barrier. In this case, the fibroblasts closest to the nest are observed to be blebbing. In addition, the green collagen bundles are entwinned with the stromal and tumor cells.

Movie S3. Related to Figure 6

The 3D FIB-SEM volume collected at 4 nm/voxel resolution from Bx2 shows ultrastructural features at the nanoscale in a 25 x 20 x 10 μm^3^ volume. Similar to the Bx1 volume, aberrant nuclear morphology, clustered macropinosomes, organized mitochondria, and the presence of lysosomes are all observed. However, the cell-cell interactions are remarkable. The center tumor cell squeezing between the surrounding cells is observed to have micron-long protrusions, while its neighbors have clear lamellipodia.

Movie S4. Related to Figure 6

Live cell stochastic optical reconstruction microscopy (STORM) imaging of HER2 in a SKBR3 (HER2+ breast cancer) cell immediately after EGF treatment (at 10 ng/mL), highlighting the dynamics of HER2-enriched FLPs during a ∼4-minute period. Note that the tips of most FLPs remained at their original locations and the cell body extended significantly in the form of lamellipodia toward the tips of the FLPs.

